# Presynaptic α_2_δ subunits are key organizers of glutamatergic synapses

**DOI:** 10.1101/826016

**Authors:** Clemens L. Schöpf, Stefanie Geisler, Ruslan I. Stanika, Marta Campiglio, Walter A. Kaufmann, Benedikt Nimmervoll, Bettina Schlick, Ryuichi Shigemoto, Gerald J. Obermair

**Author notes:** Address correspondence to: Gerald J. Obermair, Univ.-Prof. Dr.

## Abstract

In nerve cells the genes encoding for α_2_δ subunits of voltage-gated calcium channels (VGCCs) have been linked to synaptic functions and neurological disease. Here we show that α_2_δ subunits are essential for the formation and organization of glutamatergic synapses. Using a cellular α_2_δ subunit triple loss-of-function model, we demonstrate a failure in presynaptic differentiation associated with the downscaling of postsynaptic AMPA receptors and the postsynaptic density. The role of α_2_δ isoforms as synaptic organizers is highly redundant, as each individual α_2_δ isoform can rescue presynaptic calcium channel trafficking and expression of synaptic proteins. Mutating the MIDAS site in α_2_δ-2 dissociates rescuing presynaptic synapsin expression from calcium channel trafficking, suggesting that the regulatory role of α_2_δ subunits is independent from its role as a calcium channel subunit. Our findings influence the current view on excitatory synapse formation. Firstly, our study suggests that postsynaptic differentiation is secondary to presynaptic differentiation. Secondly, the dependence of presynaptic differentiation on α_2_δ implicates α_2_δ subunits as potential nucleation points for the organization of synapses. Finally, our results suggest that α_2_δ subunits act as trans-synaptic organizers of glutamatergic synapses, thereby aligning the synaptic active zone with the postsynaptic density.

---

In synapses neurotransmitter release is triggered by the entry of calcium through voltage-gated calcium channels (VGCCs). Neuronal VGCCs consist of an ion-conducting α_1_ subunit and the auxiliary β and α_2_δ subunits. α_2_δ subunits, the targets of the widely prescribed anti-epileptic and anti-allodynic drugs gabapentin and pregabalin, are membrane-anchored extracellular glycoproteins, which modulate VGCC trafficking and calcium currents (Arikkath and Campbell, 2003; Dolphin, 2013; Geisler et al., 2015; Obermair et al., 2008; Zamponi et al., 2015). In nerve cells α_2_δ subunits have been linked to neuropathic pain and epilepsy (Zamponi et al., 2015), they interact with mutant prion proteins (Senatore et al., 2012) and have been proposed to regulate synaptic release probability (Hoppa et al., 2012). Importantly, all α_2_δ isoforms are implicated in synaptic functions. Presynaptic effects of α_2_δ-1, for example, may be mediated by an interaction with α-neurexins (Brockhaus et al., 2018) or NMDARs (e.g. (Chen et al., 2018; Zhou et al., 2018)). In contrast, postsynaptic α_2_δ-1 acts as a receptor for thrombospondins (Eroglu et al., 2009) and promotes spinogenesis via postsynaptic Rac1 (Risher et al., 2018). α_2_δ-2 is necessary for normal structure and function of auditory hair cell synapses (Fell et al., 2016) and has been identified as a regulator of axon growth and hence a suppressor of axonal regeneration (Tedeschi et al., 2016). Importantly, a splice variant of α_2_δ-2 regulates postsynaptic GABA_A_-receptor abundance and axonal wiring (Geisler et al., 2019). In invertebrates, α_2_δ loss-of-function was associated with abnormal presynaptic development in motoneurons (Caylor et al., 2013; Kurshan et al., 2009) and in mice the loss of α_2_δ-3 results in aberrant synapse formation of auditory nerve fibers (Pirone et al., 2014). Finally, α_2_δ-4 is required for the organization of rod and cone photoreceptor synapses (Kerov et al., 2018; Wang et al., 2017).

Despite these important functions, knockout mice for α_2_δ-1 and α_2_δ-3 show only mild neurological phenotypes (Fuller-Bicer et al., 2009; Geisler et al., 2015; Landmann et al., 2019; Landmann et al., 2018; Neely et al., 2010; Zhou et al., 2018). In contrast, mutant mice for α_2_δ-2 (ducky) display impaired gait, ataxia, and epileptic seizures (Barclay et al., 2001), all phenotypes consistent with a cerebellar dysfunction due to the predominant expression of α_2_δ-2 in the cerebellum. Hence, in contrast to the specific functions of α_2_δ isoforms (see above) the phenotypes of the available knockout or mutant mouse models suggest a partial functional redundancy in central neurons. Moreover, detailed mechanistic insights into the putative synaptic functions of α_2_δ subunits are complicated by the simultaneous and strong expression of three isoforms (α_2_δ-1 to −3) in neurons of the central nervous system (Schlick et al., 2010).

In this study, by transfecting cultured hippocampal neurons from α_2_δ-2/-3 double-knockout mice with shRNA against α_2_δ-1, we developed a cellular α_2_δ subunit triple loss-of-function model. Excitatory synapses from these cultures show a severe failure of synaptic vesicle recycling associated with loss of presynaptic calcium channels and presynaptic vesicle-associated proteins as well as a reduced size of the presynaptic active zone. Lack of presynaptic α_2_δ subunits also induces a failure of postsynaptic PSD-95 and AMPA receptor localization and a thinning of the postsynaptic density (PSD). Each individual α_2_δ isoform (α_2_δ-1 to −3) could rescue the severe phenotype revealing the highly redundant role of presynaptic α_2_δ isoforms in glutamatergic synapse formation and differentiation. Together our results show that α_2_δ subunits regulate presynaptic differentiation as well as the trans-synaptic alignment of postsynaptic receptors and are thus critical organizers of glutamatergic synapses.

## Experimental Procedures

### Breeding of α_2_δ-2/-3 double-knockout mice

Double-knockout mice and littermate controls were obtained by crossbreeding double heterozygous α_2_δ-3^+/-^, α_2_δ-2^+/du^ mice, both backcrossed into a c57BL/6N background for more than 10 generations. α_2_δ-3 knockout (Cacna2d3^tm1Dgen^) strains generated by Deltagen (San Mateo, CA, USA) and ducky (Cacna2d2^du^; α_2_δ-2^du/du^) mice were obtained from The Jackson Laboratory (Bar Harbor, ME, USA). One week before delivery, male mice were separated from the breeding cages and BALB/c foster mothers were included. Mice were bred and maintained at the central laboratory and animal facility of the Medical University Innsbruck according to national and EU regulations and conforming to the Austrian guidelines on animal welfare and experimentation. Animal protocols, including breeding of single- and double-knockout mice, were approved by the Austrian Federal Ministry of Science, Research and Economy (BMWFW-66.011/0113-WF/V/3b/2014 and BMWFW-66.011/0114-WF/V/3b/2014). Survival rate of different α_2_δ double-knockout pups was continuously monitored over 3 years, whereby the guidelines for identifying humane endpoint criteria were strictly applied. The number of animals used for this project was annually reported to the Austrian Federal Ministry of Science, Research and Economy (bmwfw).

### Tattooing and genotyping of potential double-knockout mice

To identify mice for genotyping, newborn pups were tattooed on the paws using a sterile needle with green tattoo paste (ketschum.mfg.co).

### Genotyping

DNA was extracted by incubating a tail biopsy of ∼1-2 mm length in 100 µl of 25 mM NaOH at 95 °C for 30 min followed by cooling to 4°C and the addition of 100 µl of 40 mM Tris-HCl neutralization buffer. The PCR reaction buffer further contained 1.25 mM MgCl, 0.125 mM dNTP’s, 1 mM 5x Green GoTaq Flexi Buffer, 0.5 mM Green GoTaq Polymerase (Promega) and 2 µl of genomic DNA. Probes were analyzed using standard PCR conditions and gel electrophoresis. *Cacna2d3*^*tm1Dgen*^ *(α*_*2*_*δ-3 knockout mice):* The following primers were used for detecting the wildtype allele (F1-R, 183 bp fragment) and the knockout allele (F2-R, 331 bp fragment): F1:5’- TAGAAAAGATGCACTGGTCACCAGG-3’; F2: 5’-GGGCCAGCTCATTCCTCCCACTCAT-3’, R: 5’-GCAGAAGGCACATTGCCATACTCAC-3’.

### Cacna2d2^du^(α_2_δ-2^du/du^, ducky mice)

The evaluation of the ducky mutation was performed as previously described (Brodbeck et al., 2002). PCR with the primers du-F, 5’- ACCTATCAGGCAAAAGGACG-3’ and du-R 5’- AGGGATGGTGATTGGTTGGA-3’ revealed a fragment of 541 bp for both the wildtype and the du allele from a region that is duplicated in the du allele. Digestion with BspHI (New England Biolabs) resulted in two fragments of 286 bp and 273 bp for the du allele, whereas the fragment from the wild-type allele remained uncut. Wildtype mice could be identified by the presence of a single band upon agarose gel electrophoresis. Heterozygous and homozygous (du/du) mice each showed two bands and were preliminarily distinguished based on the relative intensity of the double band.

### Quantitative TaqMan copy number RT-PCR

In order to ultimately confirm the genomic duplication of the Cacna2d2 gene in α_2_δ-2^du/du^ mice we developed a custom designed copy number (CN) qPCR assay. Tissue biopsies of putative knockout and littermate controls were incubated at 55°C overnight constantly shaking at 550 rpm, using 250 µl Direct PCR Tail Lysis reagent (PeqLab) and 2.5 µl Protease K (20mg/ml, Roche). Following incubation, Protease K was inactivated by incubation at 85°C for 45 min constantly shaking at 550 rpm and subsequently DNA content was measured using a NanoDrop 2000 Spectrophotometer (Thermo Scientific). For each reaction 8 µl DNA (5 ng/µl) together with 10 µl TaqMan Mastermix, 1 µl Cacna2d2 CN assay labeled with a FAM dye (assay ID: Mm00270662-cn), and 1 µl transferrin receptor (Tfrc) assay as reference gene containing a VIC dye (catalogue number 4458366) were used. Chemicals were purchased from Thermo Fisher Scientific and samples were analyzed in triplicates with a 7500 fast real time PCR machine (Thermo Fisher Scientific). Relative gene expression was calculated by using the ^ΔΔ^C_T_-method (Schmittgen and Livak, 2008) normalized to wildtype control samples yielding ratios of 1 for wildtype samples (2 alleles), 1.5 for heterozygous samples (3 alleles), and 2 for homozygous ducky samples (4 alleles) (see Suppl.Fig. 2).

The abundance of different α_2_δ subunit transcripts in cDNA from cultured hippocampal neurons or hippocampus tissue was assessed by TaqMan quantitative PCR (qPCR) using a standard curve method as previously described (Schlick et al., 2010). TaqMan gene expression assays specific for the four α_2_δ isoforms were designed to span exon–exon boundaries and purchased from Applied Biosystems. The following assays were used [name (gene symbol), assay ID (Applied Biosystems)]: α_2_δ-1 (Cacna2d1), Mm00486607_m1; α_2_δ-2 (Cacna2d2), Mm00457825_m1; α_2_δ-3 (Cacna2d3), Mm00486613_m1; α_2_δ-4 (Cacna2d4), Mm01190105_m1. Expression of hypoxanthine phosphoribosyl-transferase 1 (HPRT1; Mm00446968_m1) was used as endogenous control. The qPCR (50 cycles) was performed in duplicates using total cDNA and the specific TaqMan gene expression assay for each 20 µl reaction in TaqMan Universal PCR Master Mix (Applied Biosystems). Analyses were performed using the 7500 Fast System (Thermo Fisher Scientific). The Ct values for each gene expression assay were recorded in each sample and molecule numbers were calculated from the respective standard curve (Schlick et al., 2010). Expression of HPRT1 was used to evaluate the total mRNA abundance and for normalization to allow a direct comparison between the expression levels in the different genotypes.

### Primary cultured hippocampal neurons

Low-density cultures of hippocampal neurons were prepared from putative P0-P3 du/α_2_δ-3 double-knockout mice and littermate controls as described previously (Di Biase et al., 2011; Kaech and Banker, 2006; Obermair et al., 2003; Obermair et al., 2004). Briefly, dissected hippocampi were dissociated by trypsin treatment and trituration. For imaging experiments neurons were plated on poly-L-lysine-coated glass coverslips in 60 mm culture dishes at a density of ∼3500 cells/cm^2^. After plating, cells attached for 3-4 h before transferring the coverslips neuron-side down into a 60 mm culture dish with a glial feeder layer. For electrophysiology neurons were plated directly on top of glial cells as previously reported (Stanika et al., 2012). Neurons and glial feeder layer were maintained in serum-free neurobasal medium supplemented with Glutamax and B-27 (all from Invitrogen). Ara-C (5μM) was added 3d after plating and, once a week, 1/3 of the medium was removed and replaced with fresh maintenance medium.

### Primary co-cultures of striatal and cortical neurons and transfection procedure

Co-cultures of GABAergic striatal medium spiny neurons (MSNs) and glutamatergic cortical neurons were prepared from P0-P3 du/α_2_δ-3 double-knockout mice and littermate controls (α_2_δ-3 knockout) as described previously (Geisler et al., 2019). Briefly, striatal and cortical tissue of each pup was separately collected and dissociated by trypsin treatment and trituration. Subsequently, expression plasmids were introduced into MSNs using Lipofectamine 2000-mediated transfection (Invitrogen) as described previously (Obermair et al., 2004). To this end, ∼2.4×10^5^ striatal neurons were transfected for 20 min in a 37°C water bath keeping the total volume to ≥ 1 ml with NBKO. Triple-knockout MSNs were generated by employing pβA-eGFP-U6-α_2_δ-1-shRNA (Obermair et al., 2005) knock-down in du/α_2_δ-3 double-knockout neurons. Littermate controls were transfected with pβA-eGFP, yielding α_2_δ-3 knockout neurons in which α_2_δ-1 and α_2_δ-2 are thus still present. After 20 min the cell suspension was directly seeded on poly-L-lysine coated glass coverslips within a 60 mm culture dish containing 4 ml of pre-warmed NPM and striatal neurons were allowed to attach at 37°C. For the entire transfection procedure, dissociated cortical neurons were maintained in HBSS in a 15 ml tube at 37°C and occasionally swirled. After 2 h transfection of striatal neurons was stopped by replacing the transfection-plating solution with 5 ml of fresh, pre-warmed NPM and untransfected cortical neurons were seeded onto striatal neurons in a ratio of 2 (cortical neurons) to 3 (MSNs) at a total density of ∼14,000 cells/cm^2^. Subsequently, cortical cells were allowed to attach for 3–4 h until coverslips were transferred neuron-side down into a 60 mm culture dish containing a glial feeder layer. Ara-C treatment and maintenance of neurons and glia was done as described above. Cells were processed for immunostaining at 22-24 DIV.

### Transfection of hippocampal neurons

Expression plasmids were introduced into neurons at 6 days in vitro using Lipofectamine 2000-mediated transfection (Invitrogen) as described previously (Obermair et al., 2004). Triple-knockout cultures were established by employing pβA-eGFP-U6-α_2_δ-1-shRNA (Obermair et al., 2005) knock-down in du/α_2_δ-3 double-knockout neurons. Littermate controls were transfected with pβA-eGFP. For co-transfection/rescue experiments (pβA-eGFP-U6-α_2_δ-1-shRNA plus pβA-α_2_δ-2 or pβA-α_2_δ-3) 1.5 μg of total DNA was used at a molar ratio of 1:2, respectively. Cells were processed for patch clamp experiments and immunostaining/FM-dye loading at 14-16 DIV and 17-25 DIV, respectively after plating.

### Molecular biology

To facilitate neuronal expression all constructs were cloned into a eukaryotic expression plasmid containing a neuronal chicken β-actin promoter, pβA (Fischer et al., 1998). Cloning of all constructs was confirmed by sequence (Eurofins Genomics, Germany) and sequences were deposited in Genbank.

pβA-α_2_δ-1-v2: Mouse α_2_δ-1 was cloned from genomic cDNA derived from mouse cerebellum. Primer sequences were selected according to Genbank NM-001110844. Briefly, the cDNA of α_2_δ-1 was amplified by PCR in three fragments. The forward primer used for amplifying fragment 1 introduced a NotI site and the Kozak sequence (CCTACC) upstream the starting codon and the reverse primer used for amplifying fragment 3 introduced a KpnI and a SalI site after the stop codon. Fragment 2 (nt 1442-2564) was MfeI/BamHI digested and fragment 3 (nt 2335-3276) was KpnI/BamHI digested and co-ligated in the corresponding MfeI/KpnI sites of the pβA vector, yielding an intermediate construct. Fragment 1 (nt 1-1575) was NotI/MfeI digested and co-ligated with the SalI/MfeI digested intermediate construct, containing fragment 2 and 3, and the NotI/SalI digested pβA vector, yielding pβA-α_2_δ-1-v2 (Genbank accession number MK327276; (Geisler et al., 2019)).

pβA-2HA-α_2_δ-1-v2: The putative signal peptide (aa1-24) was predicted using Signal P (SignalP 4.0: discriminating signal peptides from transmembrane regions) (Petersen et al., 2011). A double HA tag (2HA) followed by a TEV cleavage site was introduced between the third and fourth amino acids after the predicted signal peptide cleavage site of mouse α_2_δ-1, i.e. residue F27. Introduction of this sequence did not alter the predicted cleavage site. Briefly the cDNA sequence of α_2_δ-1 (nt 1–516) was PCR amplified with overlapping primers introducing the double HA tag and the TEV cleavage site in separate PCR reactions using pβA-α_2_δ-1 as template. The two separate PCR products were then used as templates for a final PCR reaction with flanking primers to connect the nucleotide sequences. The resulting fragment was then NotI/BglII digested and ligated into the corresponding sites of pβA-α_2_δ-1, yielding pβA-2HA-α_2_δ-1-v2 (Geisler et al., 2019).

pβA-SEP-α_2_δ-1: α_2_δ-1 (Genbank accession No. M21948 (Ellis et al., 1988)) was cloned into pβA and the GFP variant super-ecliptic pHluorin (SEP) was inserted after the signal sequence (after aa 26) yielding pβA-SEP-α_2_δ-1.

pβA-α_2_δ-2-v1: Mouse α_2_δ-2 was cloned from genomic cDNA from mouse brain. Primer sequences were selected according to Genebank NM-001174047. The cDNA of α_2_δ-2 was amplified by PCR in 4 fragments. The forward primer used for amplifying fragment 1 introduced a HindIII site and the Kozak sequence (CCTACC). Fragment 1 was isolated from cerebellum, while the other three fragments from hippocampus. Fragment 1 (nt 1-686) and Fragment 2 (nt 323-1294) were HindIII/BamHI and BamHI/EcoRI digested respectively, and co-ligated in the corresponding HindIII/EcoRI sites of the pBS (Bluescript) vector, yielding the intermediate construct pBS-α_2_δ-2-part1. Fragment 3 (nt 1137-2359) was EcoRI/PmlI digested and ligated into the corresponding sites of the pSPORT vector, yielding the intermediate construct pSPORT-α_2_δ-2-part2. Fragment 4 (nt 2226-3444) was BmtI/XbaI digested and ligated into the corresponding sites of pSPORT-part2, yielding the intermediate construct pSPORT-α_2_δ-2-part3. pSPORT-α_2_δ-2-part3 was EcoRI/XbaI digested and the band containing fragments 3-4 (bp 1137-3444) was ligated into pBS-α_2_δ-2-part1, yielding pBS-α_2_δ-2. This construct was then HindIII/XbaI digested and the cDNA of α_2_δ-2 was ligated into the pβA vector, yielding pβA-α_2_δ-2-v1 (Genbank accession number MK327277; (Geisler et al., 2019)).

pβA-2HA-α_2_δ-2-v1: A putative signal peptide was not reliably predicted using Signal P (SignalP 4.0: discriminating signal peptides from transmembrane regions), (Petersen et al., 2011) however the highest prediction showed that the signal peptide comprises residues 1-64. The 2HA tag followed by a thrombin cleavage site was therefore introduced after the predicted signal peptide cleavage site of mouse α_2_δ-2, i.e. residue A64. Introduction of this sequence did not alter the predicted cleavage site. Briefly the cDNA sequence of α_2_δ-2 (nt 1–761) was PCR amplified with overlapping primers introducing the double HA tag and the thrombin cleavage site in separate PCR reactions using pβA-α_2_δ-2 as template. The two separate PCR products were then used as templates for a final PCR reaction with flanking primers to connect the nucleotide sequences. The resulting fragment was then HindIII/AflII digested and ligated into the corresponding sites of pβA-α_2_δ-2, yielding pβA-2HA-α_2_δ-2-1 (Geisler et al., 2019).

pβA-2HA-α_2_δ-2-ΔMIDAS: The three divalent metal-coordinating amino acids (D300, S302, and S302) of the MIDAS domain of α2δ-2 were mutated to alanines by SOE-PCR. Briefly the cDNA sequence of 2HA-α_2_δ-2 (nt 1–1369) was PCR amplified with overlapping primers mutating the three amino acids into alanines in separate PCR reactions using pβA-2HA-α_2_δ-2 as template. The two separate PCR products were then used as templates for a final PCR reaction with flanking primers to connect the nucleotide sequences. The resulting fragment was then HindIII/EcoRI digested and co-ligated with HindII/XbaI and XbaI/EcoRI fragments of pβA-2HA-α_2_δ-2, yielding pβA-2HA-α_2_δ-2-ΔMIDAS.

pβA-α_2_δ-3: Mouse α_2_δ-3 was cloned from genomic cDNA from mouse hippocampus. Primer sequences were selected according to Genebank NM-009785. Briefly, the cDNA of α_2_δ-3 was amplified by PCR in four fragments. The forward primer used for amplifying fragment 1 introduced a NotI site and the Kozak sequence (CCTACC) upstream the starting codon. Fragment 3 (nt 1520-2817) was then SacI/PstI digested and fragment 4 (nt 2727-3276) was DraI/PstI digested and coligated in the corresponding SacI/SmaI sites of the pSPORT vector, yielding an intermediate construct. Fragment1 (nt 1-653) was then NotI/BamHI digested and fragment2 (535-1636) was then SacI/BamHI digested and co-ligated with the SacI/NotI digested intermediate construct, containing fragment3 and 4, yielding pSPORT-α_2_δ-3. The cloned cDNA of α_2_δ-3 was then NotI/RsrII digested and ligated into the corresponding sites of the pβA vector, yielding pβA-α_2_δ-3 (Genbank accession number MK327280; (Geisler et al., 2019)).

pβA-2HA-α_2_δ-3: The putative signal peptide (aa 1-28) was predicted using Signal P (SignalP 4.0: discriminating signal peptides from transmembrane regions) (Petersen et al., 2011). The 2HA tag followed by a thrombin cleavage site was therefore introduced after the predicted signal peptide cleavage site of mouse α_2_δ-3, i.e. residue D28. Introduction of this sequence did not alter the predicted cleavage site. Briefly the cDNA sequence of α_2_δ-3 (nt 1–653) was PCR amplified with overlapping primers introducing the double HA tag and the thrombin cleavage site in separate PCR reactions using pβA-α_2_δ-3 as template. The two separate PCR products were then used as templates for a final PCR reaction with flanking primers to connect the nucleotide sequences. The resulting fragment was then NotI/BsrGI digested and ligated into the corresponding sites of pβA-α_2_δ-3, yielding pβA-2HA-α_2_δ-3 (Geisler et al., 2019).

pβA-eGFP-U6-shRNA-α_2_δ-1: In brief siRNA target sequences corresponding to the α_2_δ-1 coding region (Cacna2d1, GenBankTM accession number M_009784, see (Obermair et al., 2005)) were selected and tested for efficient knockdown. The siRNA was expressed as shRNA under the control of a U6 promoter (derived from the pSilencer1.0-U6 siRNA expression vector, Ambion Ltd., Huntington, Cambridgeshire, UK) cloned into pβA-eGFP plasmid. For lentiviral expression α_2_δ-1 shRNA was cloned into pHR as previously described (Subramanyam et al., 2009).

### Electrophysiology

Calcium channel activity was recorded using the whole-cell patch-clamp technique as described previously (Stanika et al., 2012) with modifications. Patch pipettes were pulled from borosilicate glass (Harvard Apparatus) and had resistances of 2.5–4 MΩ when filled with the following (in mM): 120 cesium methanesulfonate, 1 MgCl_2_, 0.1 CaCl2, 10 HEPES, 0.5 EGTA, 4 Mg-ATP, 0.3 Na-GTP (pH 7.2 with CsOH). The bath solution contained the following (in mM): 10 BaCl2, 110 NaCl, 20 TEA-Cl, 5 4-AP, 10 HEPES, 2 MgCl2, 3 KCl, 10 Glucose, 0.001 TTX (pH 7.4 with NaOH). Currents were recorded with an EPC 10 amplifier controlled by Patch Master Software (HEKA Elektronik Dr. Schulze GmbH, Germany). Linear leak and capacitive currents were digitally subtracted with a P/4 prepulse protocol. The current–voltage dependence was fitted according to a Boltzmann equation: I=G_max_·(V-V_rev_) / (1+exp((V-V1/2) / k)) where G_max_ is the maximum conductance of endogenous calcium channels, V_rev_ is the extrapolated reversal potential of the calcium current, V_1/2_ is the potential for half-maximal conductance, and k is the slope. Cells were depolarized from a holding potential of - 70 mV to +60 mV with 10 mV steps, 10mM Barium was used as a charge carrier.

### FM-dye loading

Live cell cultured hippocampal neurons at DIV 17-25 were pre-incubated in 2.5mM KCl Tyrode solution containing (130 mM NaCl, 2.5 mM KCl, 2 mM CaCl_2,_ 2 mM MgCl_2_*6H_2_O, 10 mM HEPES, 30 mM glucose, pH 7.4) in a specialized Ludin-chamber (Life Imaging services, CH-4057 Basel Switzerland) (Nimmervoll et al., 2013). To block network activity 10 μM CNQX and 50 μM AP5 (both Tocris Bioscience, Bristol, UK) was present in all solutions and the temperature was kept at 37°C. Cells were loaded with FM4-64 dyes upon 60 mM KCl depolarization followed by a continuous washout with Tyrode solution (2.5 mM) using an inverted Axiovert 200 M setup (Carl Zeiss Light Microscopy, Göttingen, Germany) connected to a Valve Link perfusion system. For quantification, FM4-64 and eGFP images were matched. For analysis presynaptic varicosities of eGFP control and triple-knockout axons were selected that formed boutons along neighbouring non-transfected dendrites. Additionally phase contrast images were taken in order to monitor overall cell morphology. Average fluorescent intensities of single boutons were quantified using Metavue software.

### Immunocytochemistry

Immunolabeling of permeabilized neurons was performed as previously described (Obermair et al., 2010). Briefly, neurons were fixed in pF (4% paraformaldehyde, 4% sucrose) in PBS at room temperature. Fixed neurons were incubated in 5% normal goat serum in PBS/BSA/Triton (PBS containing 0.2% BSA and 0.2% Triton X-100) for 30 min. Primary antibodies were applied in PBS/BSA/Triton overnight at 4°C and detected by fluorochrome-conjugated secondary antibodies (Invitrogen). For staining of surface-expressed HA-tagged α_2_δ constructs, living neurons were incubated with the rat anti-HA antibody (1:100) for 30 min at 37 °C; coverslips were rinsed in HBSS and fixed in pF for 10 min. After fixation, neurons were washed with PBS for 30 min, blocked with 5% goat serum for 30 min, and labeled with anti-rat Alexa Fluor 594 (1:4000, 1h). Coverslips were mounted in p-phenylenediamine glycerol to retard photobleaching (Flucher et al., 1993) and observed with an Axio Imager microscope (Carl Zeiss) using 63×, 1.4 NA oil-immersion objective lens or with an Olympus BX53 microscope (Olympus, Tokio, Japan) using a 60× 1.42 NA oil-immersion objective lens. Images were recorded with cooled CCD cameras (SPOT Imaging Solutions, Sterling Heights, MI USA and XM10, Olympus, Tokio, Japan).

### Electron microscopy, structural analysis

Cultures of neurons were prepared as described above with the exception, that neurons were grown on coverglasses coated with a carbon layer as previously described (Campiglio et al., 2018), and fixed with 2% glutaraldehyde (Agar Scientific Ltd., Stansted, UK) in phosphate buffer (PB; 0.1 M, pH 7.4). After washing in PB three times 5 min at RT, neurons were post-fixed and stained with 0.2% osmium tetroxide (Electron Microscopy Sciences, Hatfield, PA) in PB (w/v) for 30 min at RT. After stopping reaction in PB, samples were washed in water four times 5 min at RT and stained with 0.25% uranyl-acetate (AL-Labortechnik e.U., Amstetten, Austria) in water (w/v) overnight at 4°C. They were then dehydrated in graded ethanols, infiltrated with anhydrous acetone (Merck KGaA, Darmstadt, Germany), and embedded in Durcupan™ ACM resin (Fluka, Buchs, Switzerland) using propylene oxide (Sigma) as intermedium. For polymerization, BEEM capsules (Science Services, Munich, Germany) were filled with freshly prepared Durcupan™, inverted and placed onto the neuron cultures and cured for 48 h at 60 °C. Thereafter, coverslips were removed from the block surface with the neurons remaining in the block. Serial ultrathin sections (40 and 70 nm, respectively) were cut with an ultramicrotome UC7 (Leica Microsystems). Sections were collected onto Formvar-coated copper slot grids and stained with 1% aqueous uranyl acetate and 0.3% Reynold’s lead citrate. They were examined in a Philips Tecnai 10 transmission electron microscope (TEM; Thermo Fisher Scientific GmbH) at 80 kV, equipped with a side-mounted camera MegaView III G3 (Electron Microscopy Soft Imaging Solutions [EMSIS] GmbH; Muenster, Germany). Images were processed with Radius software (EMSIS) and Photoshop (Adobe®) without changing any specific feature.

### Pre-embedding immunoelectron microscopy

Cultures of neurons were prepared as described above and fixed with 4% formaldehyde, 0.05% glutaraldehyde and 15% of a saturated picric acid solution (Sigma) in PB for 10 minutes at room temperature. To increase penetration of reagents, fixed neurons were infiltrated with increasing gradients of sucrose (5, 10 and 20%) in PB (w/v) 1h each at 4°C, flash-frozen on liquid nitrogen and rapidly thawed in lukewarm PB. Cells were then washed in Tris-buffered saline (TBS; 0.05 M, 0.9% NaCl, pH 7.4) and incubated in 10% normal goat serum (v/v) plus 2% BSA (w/v) in TBS for 1h at RT for blocking of nonspecific binding sites. Rabbit anti-GFP primary antibodies (Abcam, Cambridge, UK) were then applied in TBS plus 2% BSA at a concentration of 0.125 µg/ml overnight at 4 °C. Cells were washed four times 5 min at RT and incubated with 1.4 nm nanogold conjugated secondary antibodies (Nanoprobes, Yaphank, NY) in TBS plus 2% BSA at a concentration of 0.4 µg/ml overnight at 4 °C. Cells were washed in TBS four times 5 min at RT and post-fixed with 1% glutaraldehyde in TBS for 20 min at RT. After thorough wash in water, gold particles were silver-amplified using an HQ Silver™ enhancement kit (Nanoprobes). Samples were then processed directly for Durcupan™ embedding as described above. Alternatively, fixed neurons were treated with 0.1% Triton X-100 in TBS (T-TBS) for 20 min at RT for increasing penetration of reagents instead of the freeze-thaw procedure described above. Consequently, primary antibodies and nanogold-conjugated secondary antibodies were also diluted in T-TBS. All other conditions as antibody concentrations, infiltration times and washing steps stood the same.

### Antibodies

Primary antibodies were used as follows: rb-polyclonal anti-synapsin (1:20.000, 1:500 in combination with A350), m-monoclonal anti-synapsin-1, clone 46.1 (1:2.000, 1:500 in combination with A350), rb-polyclonal anti-Ca_v_2.1 (1:2.000) and rb-polyclonal anti-Ca_v_2.2 (1:2.000) and rb-polyclonal anti vGAT (1:2.000, 1:500 in combination with A350), all from Synaptic Systems (Göttingen, Germany). Further antibodies were m-monoclonal anti-PSD-95 (1:1.000, Affinity Bioreagent, Golden CO, USA), rat-monoclonal-anti-HA (1:1.000, Roche Diagnostics, Vienna, Austria), m-monoclonal anti GABA_A_R (1:500, Chemicon/EMD Millipore, Billerica, MA, USA) and rb-polyclonal anti-GFP (1:4000, Abcam ab6556, Cambridge, UK). Secondary antibodies used were as follows: goat anti-mouse Alexa 594 (1:4.000); goat anti-rabbit Alexa 350 (1:500), and Alexa 594 (1:4.000), and goat anti-rat Alexa 594 (1:4.000), all from Invitrogen (Fisher Scientific, Vienna, Austria); Fab’ fragments of goat anti-rabbit IgG conjugated to 1.4-nm gold particles (1:200, Nanoprobes, Yaphank, NY).

### Analysis and quantification. Synaptic expression of α_2_δ isoforms

In order to analyze the detailed synaptic localization of individual α_2_δ isoforms, each subunit was N-terminally tagged with a 2HA tag as described. A rat anti HA antibody (Invitrogen) was used to detect 2HA-tagged subunits. To quantify the distribution of eGFP, synapsin (A350) as well as 2HA-α_2_δ-1/-2/-3 (anti-HA/A594) signals, average fluorescent intensities were measured along a line through the respective synaptic bouton. Relative fluorescent intensities were processed in MS excel and finally illustrated with Photoshop CS6.

### Synaptic co-localization

In order to analyze the detailed synaptic localization of each presynaptic (Syn, Ca_V_2.1, Ca_V_2.2) as well as postsynaptic marker (PSD-95, GABA_A_-R, AMPA-in control, triple-knockout and α_2_δ-rescue conditions, linescan analysis was performed. To this end the distribution of the eGFP (A488) signal, the synapsin (A350) as well as the Ca_V_2.1, Ca_V_2.2 or PSD-95 (A594) signal, was measured along a line of 3µm cutting the respective synaptic bouton similar to the localization experiments. Average fluorescent intensities were background subtracted, plotted in MS excel and finally illustrated with Photoshop CS6.

### Single bouton quantification

Regions with spreading axons from hippocampal neurons (DIV 18-22) were selected in the eGFP channel. eGFP positive varicosities (putative synapses) were selected and the threshold was set in order to cover the entire area of individual boutons. By using the shrink region to fit tool (Metavue), each putative synapse was further measured for colocalization with synapsin (A350) or the Ca_V_2.1 (A594) fluorescent signal in the respective corresponding micrographs. In each channel-micrograph a separate background region was selected and subtracted from the average fluorescent intensities. For Ca_V_2.2, PSD-95, GABA_A_-R and AMPA-R quantification a slightly modified protocol was used because these staining patterns were, due to their subsynaptic localization, not directly co-localizing with the presynaptic corresponding eGFP signal. Therefore, the presynaptic ROI was dilated by 0.5 µm in order to avoid false positive or false negative staining patterns. In each journal it was therefore possible to measure the presynaptic marker synapsin together with Ca_V_2.1/Ca_V_2.2, PSD-95, GABA_A_-R and AMPA-R in a blinded manner. The following parameters were selected for quantification: Average fluorescent intensity/integrated fluorescent intensity/relative area of the bouton. For each neuron an average of 40-50 presynaptic varicosities were analyzed in 3-5 independent culture preparations for each condition. Single bouton quantification of GABAergic MSNs co-cultured with glutamatergic cortical neurons was done from two independent culture preparations for each condition as described (Geisler et al., 2019). Further data analysis was performed with MS Excel and Graph Pad Prism.

### Electron microscopy

Sampling areas were chosen at random from different regions of the thin sections for each sample per neuron culture and condition. Perpendicular cut excitatory spine synapses were randomly selected and micrographed within these areas. The length of presynaptic active zone and PSD was then measured for each synapse using ImageJ software (40 and 50 synapses each for unlabeled control and immunolabelled cultures, respectively). For quantifying the PSD extension, ten measurements per synapse were randomly made including the minimum and maximum extension and the mean value for each synapse was calculated (n=50 for each condition).

### Statistical analysis

Results are expressed as means ± S.E. except where otherwise indicated. Data were analyzed using a simple ANOVA with Holm-Sidak post-hoc analysis except where otherwise indicated. Data were organized and analyzed using MS Excel and Graph Pad Prism (Graph Pad Software, La Jolla, CA, USA). Graphs and figures were generated using Graph Pad Software and Adobe Phostshop CS6.

## RESULTS

### Epitope-tagged α_2_δ isoforms localize to presynaptic boutons

Three isoforms of calcium channel α_2_δ subunits are expressed in hippocampal neurons (Schlick et al., 2010), yet until today it is unclear whether all three isoforms contribute to specific neuronal and synaptic functions. A differential subcellular compartmentalization of α_2_δ isoforms could provide insights into their specific functions. Therefore, we first investigated the localization of HA-epitope-tagged α_2_δ-1, −2, and −3 in cultured hippocampal neurons. To this end a double HA-tag was engineered into N-termini of all three α_2_δ subunits cloned from mouse brain cDNA (Genbank accession numbers MK327276, MK327277, and MK327280) right after the signal sequence. Live-cell immunolabeling of the HA-epitope allows a direct and, most importantly, comparative analysis of α_2_δ isoform surface expression. Although the overall intensity of total surface expression levels differs between isoforms (α_2_δ-2 > α_2_δ-3 > α_2_δ-1), all three isoforms are localized to the somatodendritic and axonal membrane (Fig. 1a). In addition, α_2_δ-3 shows a preferential expression in the axon. However, despite these apparent overall differences all α_2_δ isoforms are expressed on the surface of axons and presynaptic membranes (Fig. 1b) suggesting that, in principle, all three isoforms can contribute to synaptic functions.

**Fig. 1.**
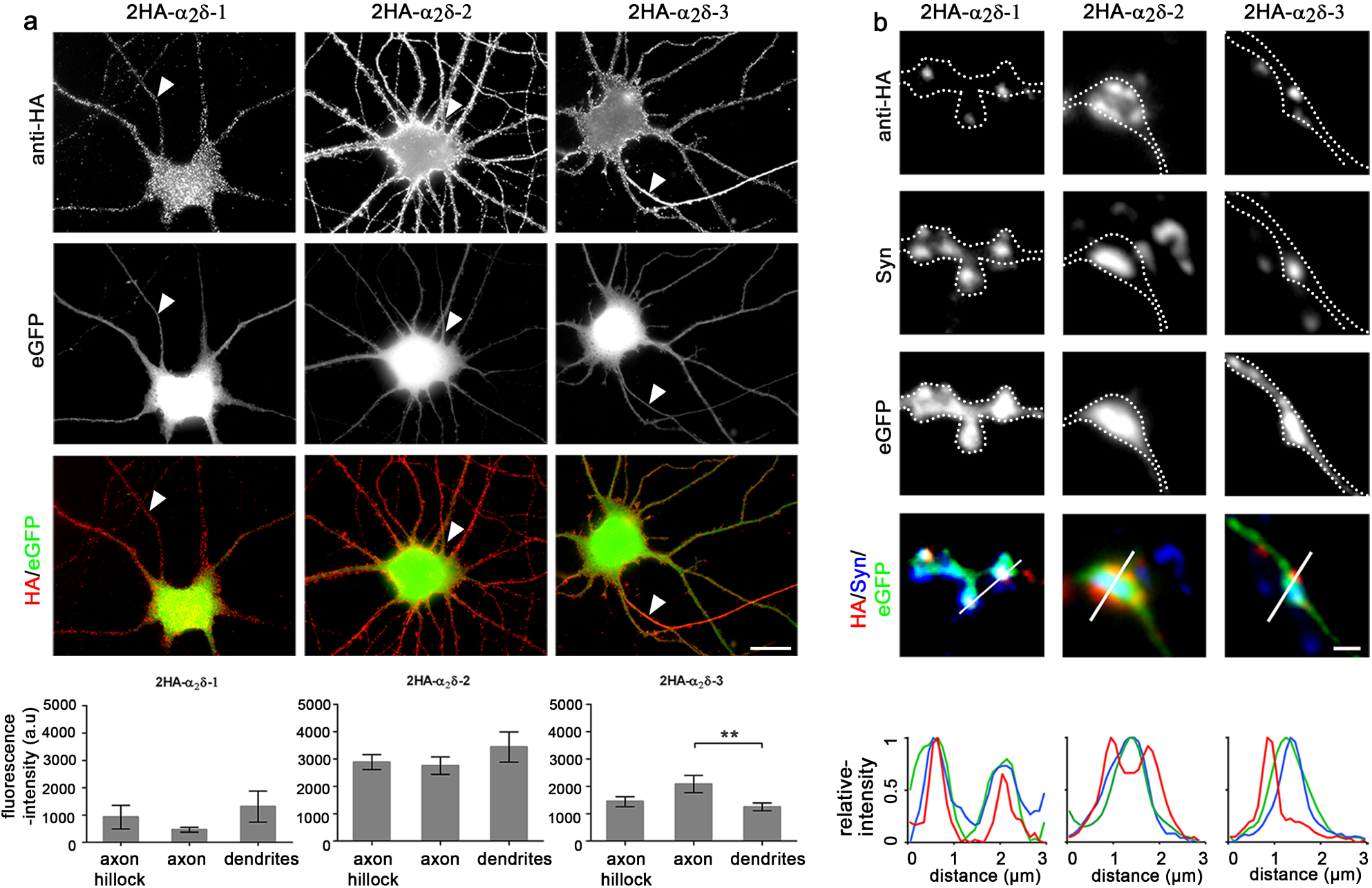
Subcellular distribution of the three neuronal α2δ subunits. Cultured hippocampal neurons were transfected with HA-tagged α_2_δ subunits together with soluble eGFP to outline the neuronal morphology and live cell labeled for the HA epitope. (**a**) Live cell staining revealed a strong expression of all three α_2_δ isoforms on the soma, dendrites and axons (arrowheads). Overall, α_2_δ-2 surface expression was up to two-fold higher when compared to α_2_δ-1 or α_2_δ-3, respectively, and α_2_δ-3 specifically accumulated in the axon. Average fluorescence intensity measurements are shown for axon hillocks, axons and dendrites. Statistics were performed with one-way ANOVA, followed by Holm-Sidak post-hoc test; α_2_δ-1: F_(2,9)_=0.02, p=0.99; α_2_δ-2: F_(2,12)_=0.83, p=0.46; α_2_δ-3: F_(2,35)_=4.0, p=0.027; **p=0.01; Error bars indicate SEM. Scale bar, 20µm. (**b**) All α_2_δ isoforms showed a synaptic localization, which is supported by the overlay or juxtaposition of the linescan peaks of synapsin (blue), HA-α_2_δ (red), and eGFP (green). α_2_δ-2 specifically accumulated in the perisynaptic membrane around the central synapsin label. Scale bar, 1µm.

### α_2_δ subunit isoforms are essential for survival

With the exception of the α_2_δ-2 mutant mouse ducky, knockout mice for α_2_δ-1 and α_2_δ-3 display only mild neuronal phenotypes, suggesting a potential and at least partial functional redundancy (see above). Therefore, in order to gain insight into the functional diversity of α_2_δ subunits, we generated double-knockout mice by pairwise cross-breeding single-knockout (α_2_δ-1, α_2_δ-3) and mutant (α_2_δ-2^du^) mice. While α_2_δ-1/-3 knockout mice are viable for up to three months similar to ducky mice, α_2_δ-1/-2 and α_2_δ-2/-3 knockout mice have a strongly reduced lifespan (Fig. 2a,b). A significant proportion of these mice require application of humane endpoints within the first postnatal week mainly due to malnutrition associated with a poor general condition. Together this shows that α_2_δ subunits serve essential functions and are necessary for survival. Moreover, the increased severity of the phenotype in double-compared to single-knockout mice also supports the idea that α_2_δ subunits act in part redundantly.

**Fig. 2.**
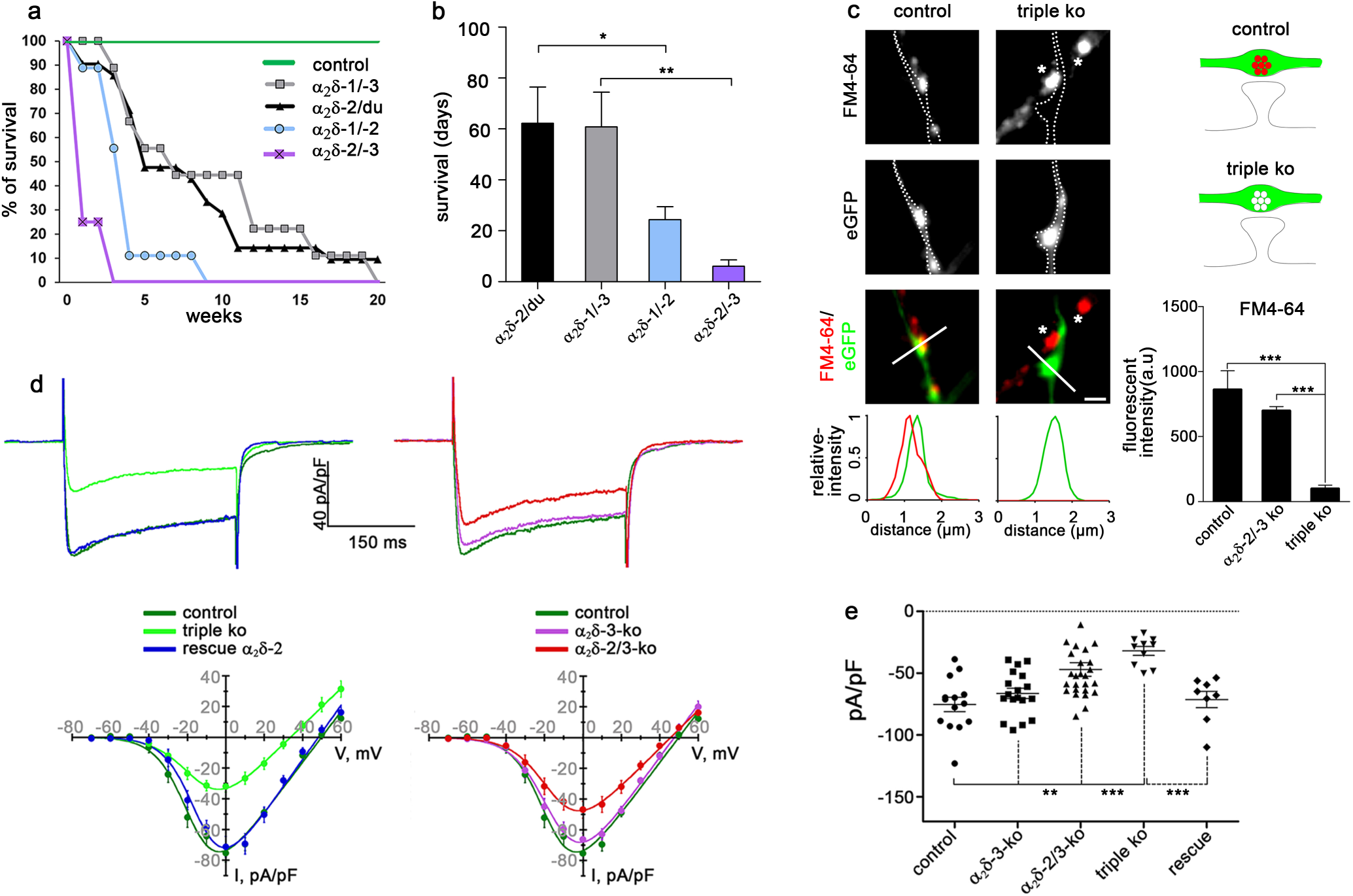
α2δ subunits are essential for survival, activity-induced synaptic recycling and normal calcium current densities. (**a**) The Kaplan Meier survival curves show an increased mortality in the distinct α_2_δ double knockout mouse models (n-numbers: 9-21). (**b**) Mean life span was significantly reduced in α_2_δ-1/-2 and α_2_δ-2/-3 double knockout mice when compared to α_2_δ-1/-3 or ducky mice (ANOVA, F_(3, 47)_=4.7, p=0.006, with Holm-Sidak post-hoc test, *p<0.05, **p<0.01). (**c**) Putative synaptic varicosities from α_2_δ triple knockout neurons failed to load FM4-64 dye upon 60 mM KCl depolarization (outline/triple ko). In contrast, control boutons transfected with eGFP only and non-transfected double knockout boutons (asterisks) showed robust uptake of the FM-dye. (ANOVA on ranks, H_(2)_= 96.6, p<0.001, with Dunn’s post-hoc test, ***p<0.001, 26-110 synapses from 1-4 culture preparations.) (**d**) Current properties of α_2_δ subunit single, double and triple knockout cultured hippocampal neurons. Representative Ba^2+^ whole-cell currents at I_max_ (upper panel) and I/V-curves (lower panel) recorded from hippocampal neurons. **left:** I/V-curves reveal a strong reduction of calcium channel currents in triple knockout neurons (triple ko), when compared to untransfected wildtype neurons or triple knockout neurons transfected with α_2_δ-2 (rescue α_2_δ-2). **right:** Current densities in α_2_δ-2/-3 double but not in α_2_δ-3 single-knockout were also reduced. For I/V curve properties see Supplementary Table 1. (**e**) Current densities at I_max_ for individual cells (ANOVA, F_(4, 71)_=11.3, p<0.001, with Holm-Sidak post-hoc test, **p<0.01, ***p<0.001, 8-26 cells from 5 culture preparations). Horizontal lines represent means, error bars SEM.

### Establishing a cellular α_2_δ-subunit triple loss-of-function model

In order to study a potential functional redundancy of α_2_δ subunits we next developed a cellular α_2_δ triple-knockout/knockdown model system by transfecting cultured hippocampal neurons from α_2_δ-2/-3 double-knockout mice with shRNA against α_2_δ-1. To this end we first confirmed efficient shRNA knockdown of α_2_δ-1 in two independent experimental settings: first, shRNA against α_2_δ-1 (Obermair et al., 2005) effectively reduced the surface expression of a heterologously expressed α_2_δ-1 isoform bearing an extracellular phluorin-tag (super-ecliptic phluorin, SEP, Suppl. Fig. 1a and b). Second, qPCR analysis of cultured hippocampal neurons virally infected with α_2_δ-1 shRNA revealed an overall 80% knockdown of α_2_δ-1 mRNA compared to untransfected (wildtype) neurons or neurons expressing scrambled control shRNA (Suppl.Fig. 1c). Considering a ∼90% infection efficiency, confirmed by eGFP expression from the same viral vector, shRNA robustly knocked down mRNA in the vast majority of infected neurons. Most importantly, shRNA knockdown of α_2_δ-1 did not affect the expression levels of the other α_2_δ isoforms (Suppl.Fig. 1c). In order to evaluate potential compensatory mechanisms we also quantified mRNA levels of all α_2_δ isoforms in hippocampal tissue from 8-week-old single knockout mice. Similar to α_2_δ-1 knockdown, neither loss of α_2_δ-2 nor of α_2_δ-3 induced compensational changes in the expression levels of the other isoforms (Suppl. Fig. 1d and e).

α_2_δ-2/-3 double knockout mice were generated by crossbreeding double heterozygous α_2_δ-2^+/du^/α_2_δ-3^+/-^ mice yielding double knockout mice at a predicted Mendelian ratio of 6.25% (Suppl. Fig. 2a). Neonatal pups (P0-2) were individually marked by paw-tattooing and genotyped for the α_2_δ-2 and α_2_δ-3 alleles (Suppl. Fig. 2b and 2c). Due to the large genomic rearrangement in ducky mice, genotyping of the ducky mutation required a confirmation employing a copy number counting qPCR approach (Suppl. Fig. 2d). Ultimately, α_2_δ triple loss-of-function hippocampal neurons were established by transfecting confirmed α_2_δ-2/-3 double knockout cultures with α_2_δ-1 shRNA and eGFP (Suppl. Fig. 2e,f).

### Failure of presynaptic differentiation in α_2_δ subunit triple loss-of-function neurons

In cultured hippocampal neurons from α_2_δ-2/-3 double knockout mice, shRNA-transfected neurons (α_2_δ-2/-3 double knockout with α_2_δ-1 shRNA knockdown, further referred to as triple-knockout) can be easily identified by the expression of soluble eGFP. Most importantly, in this experimental setting isolated axons and synaptic varicosities from transfected triple-knockout neurons can be directly compared with untransfected neighboring neurons, which still express α_2_δ-1 (Suppl. Fig. 2f). Axons from triple-knockout neurons display axonal varicosities along dendritic processes of non-transfected neighboring cells, typical to those of presynaptic boutons in cultured control neurons (compare left and right panels in Suppl. Fig. 2f). In order to test whether these boutons represent functional synapses capable of vesicle recycling we quantified the extent of depolarization-induced uptake of the styryl membrane dye FM4-64. Upon a 60mM [K^+^]-induced depolarization 68% of the axonal varicosities of triple-knockout neurons completely failed to take up FM dye and loading of the remaining 32% was strongly decreased (Fig. 2c). In contrast, neighboring untransfected (α_2_δ-1 containing) synapses (Fig. 2c, asterisks in right panel) and eGFP-transfected control neurons were readily stained with FM4-64 upon high [K^+^] treatment. This apparent failure of synaptic vesicle recycling pointed towards a severe defect in presynaptic calcium channel functions.

Indeed, voltage-clamp analysis of total somatic calcium currents identified a marked reduction of current densities by 58%, but no change in the voltage-dependence of activation of triple-knockout compared to α_2_δ-3 single-knockout or wildtype neurons (Fig. 2d,e and Suppl. Tab. 1). Notably, current densities were also strongly reduced in α_2_δ-2/-3 double-knockout neurons (by 32%, Fig. 2e), however, without a concomitant failure in FM dye uptake (confer Fig. 2c). The homologous reconstitution of α_2_δ-2 in α_2_δ triple-knockout neurons fully rescued the currents back to wildtype levels (Fig. 2d,e and Suppl. Tab. 1).

While reduced somatic calcium channel activity was to be expected in an α_2_δ-null model, the complete failure of FM dye uptake suggests a more severe failure of synaptic vesicle recycling. Therefore we next employed immunocytochemistry to test whether and to what extent the synaptic localization of presynaptic P/Q- (Ca_V_2.1) and N-type (Ca_V_2.2) calcium channels was affected (Fig. 3a,b). Strikingly, 61% and 40% of the axonal α_2_δ triple-knockout varicosities lacked detectable staining for Ca_V_2.1 and Ca_V_2.2, respectively. The remaining axonal boutons showed a strong and significant reduction of presynaptic labelling intensities (Fig. 3c,d). In agreement with defective synaptic vesicle recycling, these boutons were also deficient in synapsin staining (complete loss in 45% of the analyzed boutons, Fig. 3e). The strongly reduced presynaptic calcium channel abundance in α_2_δ triple-knockout varicosities is in line with the major role of α_2_δ subunits in enhancing calcium channel trafficking (Dolphin, 2018). However, the surprising loss of synapsin staining suggests that the lack of α_2_δ subunits also grossly affects presynaptic differentiation.

**Fig. 3.**
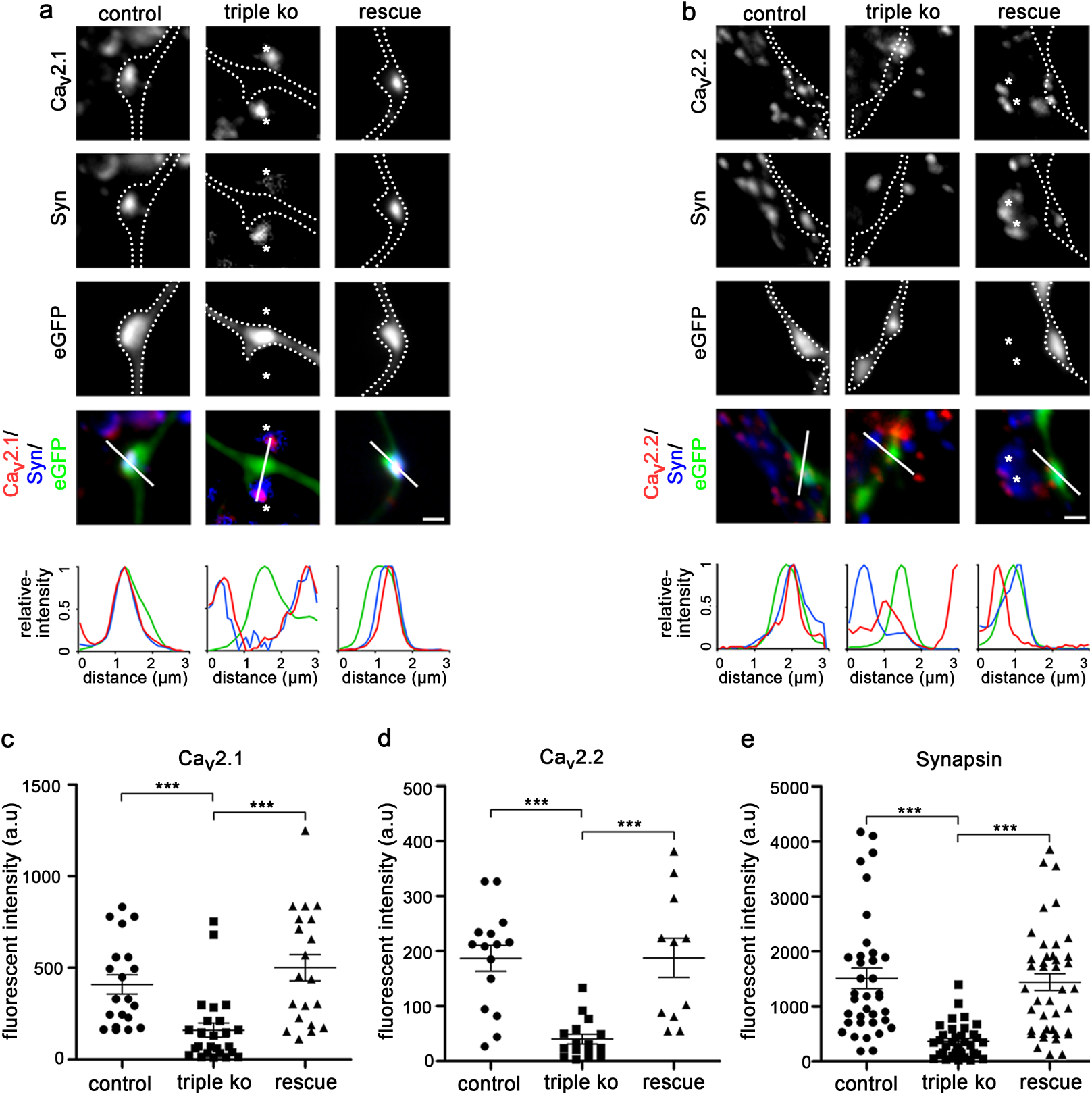
Failure of presynaptic calcium channel clustering and synapsin accumulation in α2δ subunit triple knockout neurons. (**a, b**) Immunofluoresence analysis of axonal varicosities from wildtype neurons (control, neurons transfected with eGFP only), triple knockout neurons (triple ko, α_2_δ-2/-3 double knockout neurons transfected with shRNA-α_2_δ-1 plus eGFP), and triple knockout neurons expressing α_2_δ-2 (rescue, α_2_δ-2/-3 double knockout neurons transfected with shRNA-α_2_δ-1 plus eGFP and α_2_δ-2). Putative presynaptic boutons were identified as eGFP-filled axonal varicosities along dendrites of untransfected neurons (confer Suppl. Fig. 2) and outlined with a dashed line. Immunolabeling revealed a failure in the clustering of presynaptic P/Q- (**a**, Ca_V_2.1) and N-type (**b**, Ca_V_2.2) channels as well as in the accumulation of presynaptic synapsin in varicosities from α_2_δ triple knockout neurons (middle columns). In contrast, wildtype control neurons (left columns) displayed a clear co-localization of the calcium channel clusters with synapsin in the eGFP-filled boutons. The linescan patterns recorded along the indicated line support these observations. Note that the sole expression of α_2_δ-2 (right columns) or the sole presence of α_2_δ-1 in synapses from neighboring α_2_δ-2/-3 double knockout neurons (asterisks in a and b) suffices to fully rescue presynaptic calcium channel clustering and synapsin accumulation. (**c-e**), Quantification of the fluorescence intensities of presynaptic Ca_V_2.1 (**c**), Ca_V_2.2 (**d**) and synapsin (**e**) clustering in control, triple knockout, and α_2_δ-2-expressing (rescue) triple knockout neurons (ANOVA with Holm-Sidak post-hoc test, ***p<0.001; Ca_V_2.1: F_(2, 58)_=10.8, p<0.001, 16-25 cells from 4-6 culture preparations; Ca_V_2.2: F_(2, 37)_=13.7, p<0.001, 11-16, 2-4; Synapsin: F_(2, 99)_=15.5, p<0.001, 30-36, 5-8; Horizontal lines represent means, error bars SEM). Scale bar, 1 µm.

### Presynaptic α_2_δ subunits regulate pre- and postsynaptic differentiation of excitatory glutamatergic synapses

By acting as a thrombospondin receptor, α_2_δ-1 has previously been suggested to contribute to synaptogenesis by a postsynaptic mechanism (Eroglu et al., 2009; Risher et al., 2018). Therefore, in order to distinguish between the proposed postsynaptic mechanism and the defect in presynaptic differentiation observed here, we examined triple-knockout neurons connected to neighboring non-transfected double-knockout neurons still expressing α_2_δ-1 (see Suppl. Fig. 2). In this experimental paradigm, eGFP-positive axonal processes of presynaptic triple-knockout neurons (Fig. 4a and b, left panels and sketches) can be clearly distinguished from eGFP-positive dendrites of postsynaptic triple-knockout neurons (Fig. 4a and b, right panels and sketches). These experiments demonstrate that synapse differentiation fails when the presynaptic neuron lacks all α_2_δ subunits (Fig. 4a and b, left panels). On the other hand, postsynaptic triple-knockout neurons can still form dendritic spines and receive synaptic inputs from neighboring double-knockout neurons expressing α_2_δ-1 (Fig. 4a and b, right panels). The presynaptic defect in synapse formation also induced a failure in the postsynaptic differentiation: boutons devoid of calcium channels or synapsin were either not juxtaposed to PSD-95 clusters at all (Fig. 4c) or the PSD-95 labelling was strongly reduced (Fig. 4d). Similar to the marked reduction of presynaptic synapsin and calcium channel labelling, PSD-95 was completely absent in 58% of the analyzed triple-knockout synapses. Thus, in addition to the failure in presynaptic differentiation, the lack of presynaptic α_2_δ subunits also induced a failure in postsynaptic differentiation.

**Fig. 4.**
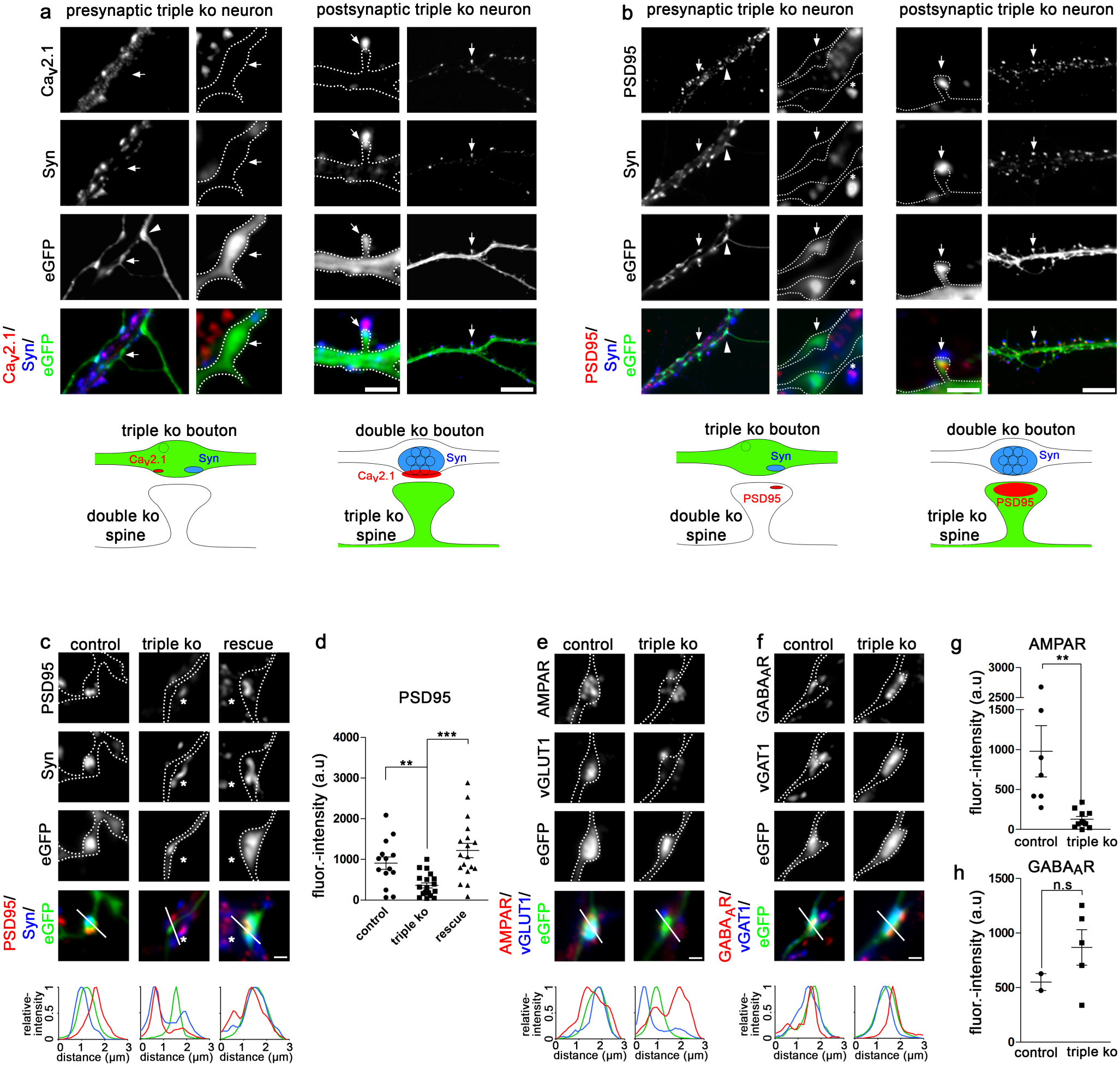
Presynaptic α2δ subunits mediate glutamatergic synapse formation and trans-synaptic differentiation. (**a, b**) Immunofluoresence micrographs of axonal varicosities from presynaptic triple knockout neurons (eGFP-positive axonal varicosities, left panels) as well as dendrites from postsynaptic triple knockout neurons (eGFP-positive dendrites, right panels). Axonal varicosities and dendrites are outlined by a dashed line; the sketches summarize the observed labeling patterns. (**a**) α_2_δ triple knockout neurons display a failure of presynaptic Ca_v_2.1 channel and synapsin clustering exclusively in presynaptic axonal varicosities (arrows and sketch, left panel). In contrast, postsynaptic triple knockout neurons developed dendritic spines opposite presynaptic boutons containing Ca_v_2.1 and synapsin clusters (arrows and sketch, right panel) formed by axons from α_2_δ-2/-3 double knockout neurons still containing α_2_δ-1. (**b**) Presynaptic α_2_δ triple knockout induces a failure of the postsynaptic PSD-95 clustering indicating a trans-synaptic action of α_2_δ subunits (arrows and sketch, left panel). Conversely, postsynaptic triple knockout neurons still receive proper synaptic input from neighboring α_2_δ-1 containing neurons as indicated by presynaptic synapsin and postsynaptic PSD-95 co-localized on triple knockout dendritic spines (arrows and sketch, right panel). Scale bars, 2 µm and 8 µm. (**c, d**) Failure of postsynaptic PSD-95 labeling opposite α_2_δ triple knockout boutons. Like the presynaptic proteins (confer Fig. 3) the sole expression of α_2_δ-2 (rescue, right column) or the sole presence of α_2_δ-1 in synapses from neighboring α_2_δ-2/-3 double knockout neurons fully rescued postsynaptic PSD-95 clustering (asterisks in c, middle column, linescans) (ANOVA, F_(2, 49)_=11.7, p<0.001, with Holm-Sidak post-hoc test, **p<0.01, ***p<0.001; 14-20 cells from 3-4 culture preparations). (**e, g**) The defect in synaptogenesis caused by loss of α_2_δ subunits specifically affects glutamatergic synapses, indicated by reduced fluorescent intensity of vGLUT1/AMPAR labeling (outline/linescan; t-test, t_(15)_=3.1, **p<0.01; 7 and 10 cells from 2 and 3 culture preparations). (**f, h**) In contrast, vGAT/GABA_A_R labeling in GABAergic synapses was not reduced in α_2_δ triple knockout neurons (outline/linescan; t-test, t_(5)_=1.16, p=0.30; 2 and 6 cells from 1 and 2 culture preparations). Error bars indicate SEM. Scale bar, 1 µm.

For analyzing whether presynaptic α_2_δ subunits are required for both excitatory and inhibitory synapse formation, we immunolabelled triple-knockout and control neurons for respective components of the presynaptic vesicle compartment and postsynaptic receptors (Fig. 4e,f). In excitatory glutamatergic neurons the lack of presynaptic staining for the vesicular glutamate transporter type 1 (vGlut1) goes along with strongly reduced clustering of postsynaptic AMPA receptors in triple-knockout synapses (Fig. 4e,g). Conversely, triple-knockout synapses from GABAergic neurons still express the presynaptic vesicular GABA transporter (vGAT) and display postsynaptic clustering of GABA_A_-receptors (Fig. 4f,h). However, it is important to note that due to the low abundance of GABAergic neurons (∼5-10% of all cultured hippocampal neurons), the extremely low availability of α_2_δ-2/-3 double knockout offspring (only 2-5 culture preparations are possible per year), and the necessity of shRNA transfection, we could only analyze two cells each for control and triple knockout conditions. Therefore, to confirm this finding we also analyzed the abundance of pre- and postsynaptic proteins in control (α_2_δ-3 knockout) and triple knockout cultured GABAergic striatal medium spiny neurons (Fig. 5). In contrast to glutamatergic hippocampal neurons, α_2_δ triple knockout in GABAergic medium spiny neurons affected neither the abundance of presynaptic vGAT (Fig. 5d) and synapsin (Fig. 5h), nor of postsynaptic GABA_A_-receptors (Fig. 5c). However, although not statistically significant, there was a tendency for reduced presynaptic Ca_V_2.1 labelling (Fig. 5g). Together this demonstrates that the severe consequence of presynaptic α_2_δ subunit triple-knockout is specific to excitatory glutamatergic neurons.

**Fig. 5.**
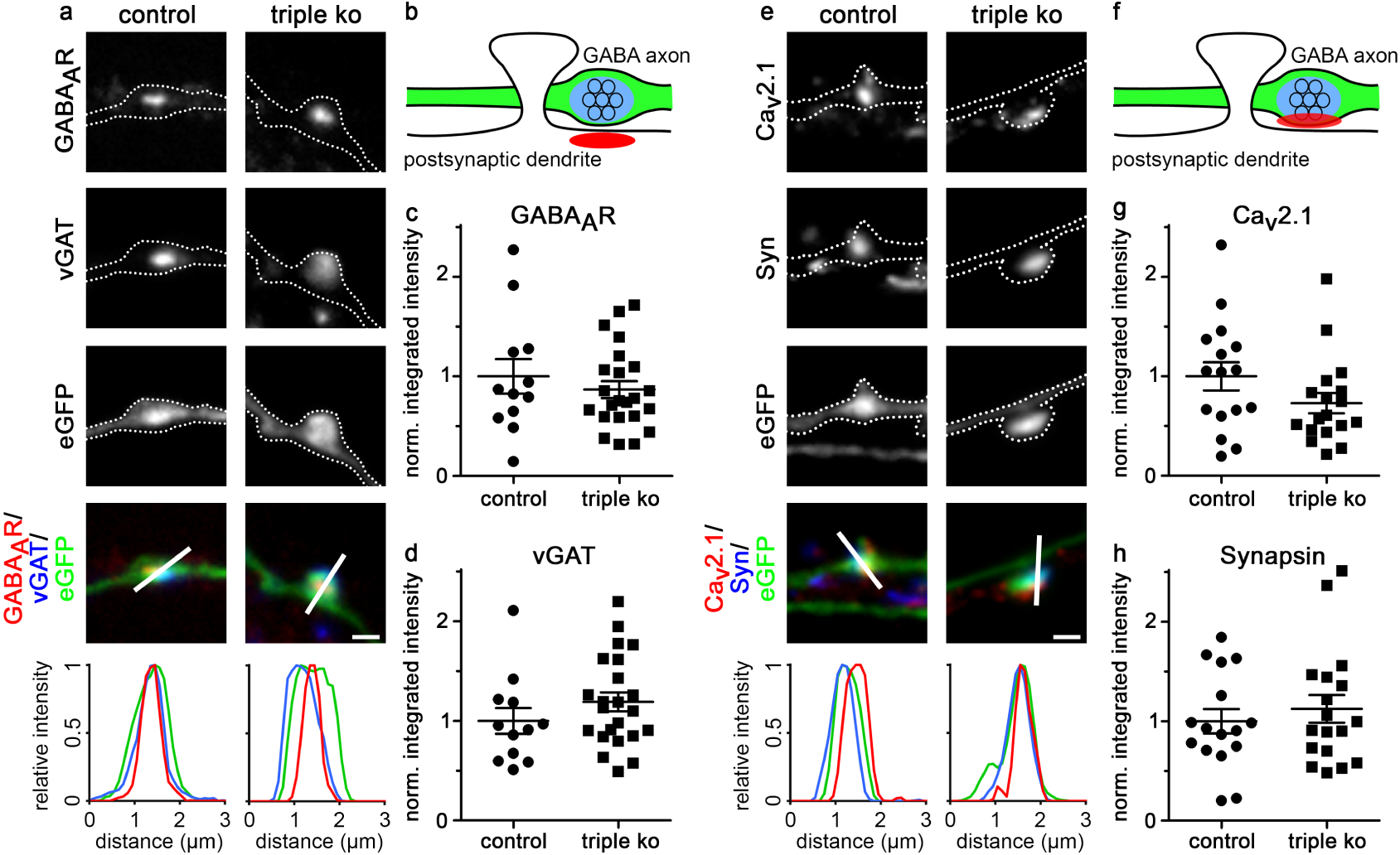
Presynaptic α2δ subunit triple knockout does not affect pre- and postsynaptic differentiation in GABAergic synapses. (**a, e**), Representative immunofluoresence micrographs of axonal varicosities from presynaptic α_2_δ-3 knockout (control) or triple knockout (triple ko) cultured GABAergic MSNs. Transfected neurons (22-24 DIV) were immunolabeled for vGAT and the GABA_A_R (**a**) and Ca_V_2.1 and synapsin (**e**). Co-localization of fluorescence signals within eGFP-filled axonal varicosities (axons are outlined with dashed lines) was analyzed using line scans. (**b, f**), Sketches depicting the expected staining patterns in (**a**) and (**e**), respectively. (**c, d, g, h**), Quantification of the respective fluorescence intensities in control and triple knockout neurons (t-test, GABA_A_R: t_(33)_=0.8, p=0.45, 12-22 cells from 2 culture preparations; vGAT: t_(33)_=-1.2, p=0.24, 12-22 cells from 2 culture preparations; Ca_V_2.1: t_(32)_=1.6, p=0.12, 16-28 cells from 2 culture preparations; synapsin: t_(32)_=-0.7, p=0.51, 16-18 cells from 2 culture preparations). Values for individual cells (dots) and means (lines) ± SEM are shown. Values were normalized to control (α_2_δ-3 knockout) within each culture preparation. Scale bar, 1 µm.

### α_2_δ subunit triple knockout affects the pre- and postsynaptic ultrastructure

Immunofluorescence labelling identified a strong reduction in the abundance of presynaptic and postsynaptic proteins in glutamatergic synapses of α_2_δ subunit triple knockout neurons. In order to test whether these presynaptic effects are associated with ultrastructural alterations we performed classical transmission electron microscopy (TEM) and pre-embedding immunoelectron microscopy. Classical TEM analysis revealed the necessity for immunolabeling shRNA-α_2_δ-1/eGFP transfected double knockout neurons in order to reliably identify the sparsely distributed triple knockout synapses weak in morphological cues. The strong immunolabeling for eGFP with the contrast intense silver-amplified gold particles, however, obscured the presynaptic ultrastructure and rendered reliable analysis of synaptic vesicle content and localization impossible. For quantifying size and extension of synaptic specializations, we first compared synapses of non-labeled wildtype control and α_2_δ-2/-3 double-knockout neurons (Fig. 6a). Analyses of 40 synapses in each condition revealed that the length of the active zone (AZ) and the PSD, the AZ/PSD ratio, as well as the PSD thickness (extension from the membrane into the cytosol) were indistinguishable between control and double-knockout neurons (mean±sem in nm, unpaired t-test; AZ: control, 433±22, double-ko, 433±20, p=0.99; PSD: control, 440±23, double-ko, 436±20, p=0.89; PSD extension: control, 28.5±1.4; double-ko, 27.2±1.1, p=0.49; AZ/PSD ratio: control, 0.986±0.004, double-ko, 0.995±0.006; p=0.19). We next performed the same analysis on eGFP-immunostained double (control eGFP) and triple knockout (triple ko) synapses (Fig. 6b). As an additional control, we measured the respective AZ and PSD parameters of non-transfected neighboring synapses (control nt), which are all double knockout for α_2_δ-2/-3. Both, AZ and PSD lengths were significantly reduced by approximately 25% in triple knockout synapses (Fig. 6c, left and middle graph), however, the AZ/PSD ratio was not altered (AZ/PSD ratio: control eGFP, 1.00±0.01, triple ko, 1.05±0.04; control nt, 0.98±0.04; ANOVA, F_(2,147)_=1.45, p=0.24). This suggests that reductions in the presynaptic AZ caused by lack of α_2_δ subunits are directly affecting the size of the PSD. Control measurements in non-transfected synapses (control nt) were indistinguishable from eGFP-transfected α_2_δ-2/-3 double knockout neurons (control eGFP). The extension of the PSD from the synaptic membrane into the cytosol was reduced by 40% in triple ko synapses compared to both controls (Fig. 6c, right graph). Taken together, these measurements reveal that presynaptic α_2_δ subunit triple knockout reduces the sizes of the presynaptic AZ and PSD as well as the thickness of the PSD.

**Fig. 6.**
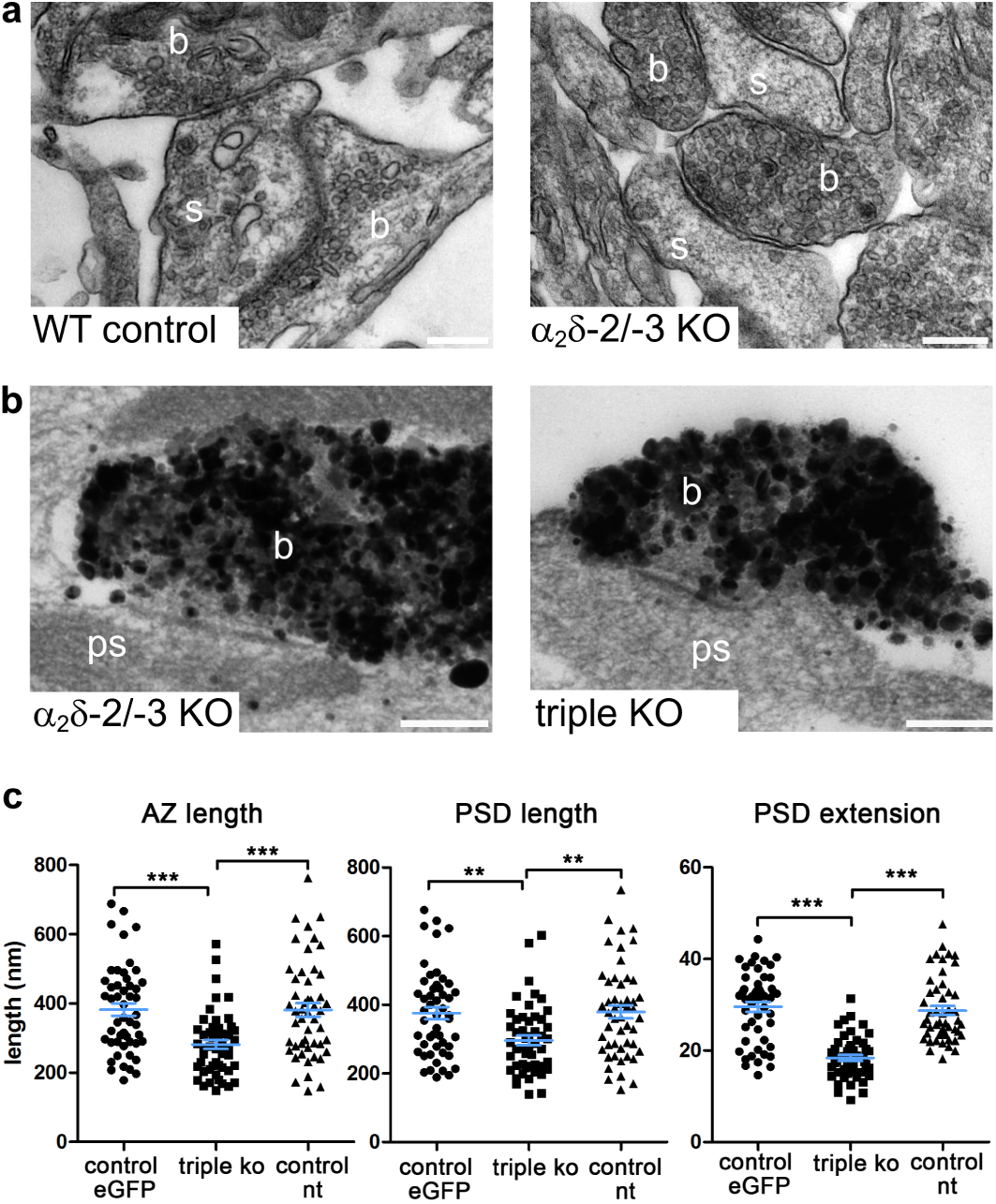
Ultrastructural analysis of pre- and postsynaptic specializations in excitatory α2δ subunit triple knockout synapses. (**a**) Exemplary EM micrographs of synaptic structures show similar presynaptic and postsynaptic differentiation in wildtype control (left) and α_2_δ-2/-3 double knockout (α_2_δ-2/-3 KO) cultured hippocampal neurons (for statistics see text). (**b**) Exemplary EM micrographs from silver-amplified eGFP-immunogold-stained presynaptic boutons and the corresponding postsynaptic region from double (α_2_δ-2/-3) and triple knockout (triple KO) synapses. (**c**) Quantitative analyses showing that both, the length of the active zone (AZ, left graph) and the postsynaptic density (PSD, middle graph) were significantly reduced in triple knockout compared to eGFP-transfected α_2_δ-2/-3 double knockout synapses in separate culture preparations (control eGFP) and non-transfected neighboring synapses within the same coverglass (control nt). In addition to the AZ and PSD length also the thickness, particularly the extension of the PSD from the membrane into the cytosol, was strongly reduced in triple knockout compared to the respective control synapses. (ANOVA with Tukey post-hoc test, **p<0.01, ***p<0.001; AZ length: F_(2, 147)_=11.3, p<0.001; PSD length: F_(2, 147)_=7.5, p<0.001; PSD extension: F_(2, 147)_=44.6, p<0.001. Horizontal lines represent means, error bars SEM). Abbreviations in EM micrographs: b, presynaptic bouton; s, dendritic spine; ps, postsynaptic compartment. Scale bars, 200 nm.

### The α_2_δ subunit triple knockout phenotype can be rescued by expression of α_2_δ-1, −2 and −3

The severe consequences of presynaptic α_2_δ triple-loss of function on pre- and postsynaptic composition and synaptic ultrastructure strongly suggests a functional redundancy. Thus, to further elucidate the potentially redundant roles of α_2_δ subunits in pre- and postsynaptic differentiation, we analyzed the propensity of each individual isoform in rescuing synapse formation and differentiation. First, α_2_δ-2/-3 double-knockout neurons, which solely express α_2_δ-1 showed a proper apposition of pre- and postsynaptic proteins (see co-localized synaptic markers near the eGFP-positive triple-knockout axons indicated by asterisks in figures 2c, 3a, b and 4b, c). Moreover, the triple-knockout phenotype could be fully rescued by the expression of both α_2_δ-2 (rescue in figures 2d,e, 3 and 4c) and α_2_δ-3 (Suppl. Fig. 3). Together this proves that the apparent critical roles of α_2_δ subunits in glutamatergic synapse formation are highly redundant between the neuronal α_2_δ isoforms.

### Expressing α_2_δ-2-ΔMIDAS in triple knockout synapses dissociates presynaptic synapsin accumulation from calcium channel trafficking

Our experiments demonstrate an essential role of α_2_δ subunits in glutamatergic synapse formation and differentiation which might be related to the failure of presynaptic calcium channel trafficking. Alternatively, however, α_2_δ subunits may act trans-synaptically and independent of the calcium channel complex, as has been previously proposed (Fell et al., 2016; Geisler et al., 2015; Geisler et al., 2019; Kurshan et al., 2009). α_2_δ subunits contain a von Willebrand factor type A (VWA) domain which, at least in α_2_δ-1 and α_2_δ-2, includes a perfect metal ion-dependent adhesion site (MIDAS). The integrity of the MIDAS motif in α_2_δ-2 is necessary for calcium current enhancement and channel trafficking (Canti et al., 2005). This finding is supported by the proposed structure of α_2_δ-1, in which the VWA domain and particularly the MIDAS is facing the surface of the pore-forming α_1_ subunit and thus predicted to be crucial for α_1_ and α_2_δ subunit interactions (Wu et al., 2016). We reasoned that mutating the MIDAS site, which has previously been shown to inhibit channel trafficking (Canti et al., 2005), may be helpful in dissociating channel-dependent from potential channel-independent functions of α_2_δ subunits. To this end we mutated the amino acids D300, S302, and S304 of α_2_δ-2 to alanines (α_2_δ-2-ΔMIDAS) and analyzed to which extent expression of α_2_δ-2-ΔMIDAS can rescue synaptic targeting of endogenous calcium channels and synapsin (Fig. 7). While α_2_δ-2-ΔMIDAS rescued presynaptic Ca_V_2.1 labelling only partially to 31% of the rescue observed with normal α_2_δ-2 (Fig. 7b), presynaptic synapsin labelling was almost fully rescued to 83% of α_2_δ-2 (Fig. 7c). Taken together, expression of α_2_δ-2-ΔMIDAS in triple knockout synapses dissociates presynaptic synapsin accumulation from calcium channel trafficking.

**Fig. 7.**
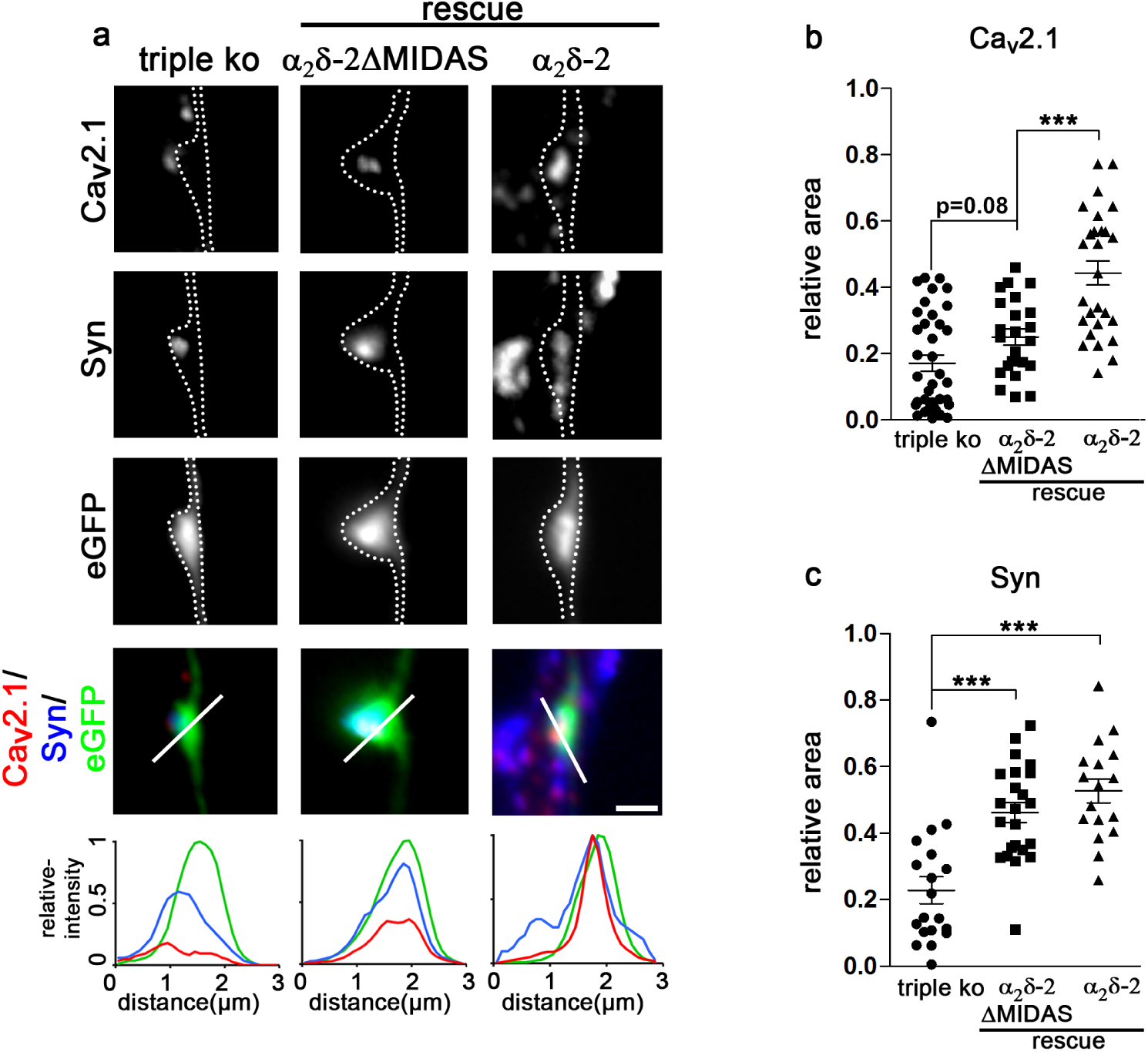
Rescuing triple-knockout synapses with α2δ-2-ΔMIDAS dissociates synapse differentiation from presynaptic calcium channel trafficking. (**a**) Immunofluoresence micrographs of axonal varicosities from presynaptic triple knockout neurons (triple ko, eGFP-positive axonal varicosities, left panels) and neurons expressing α_2_δ-2-ΔMIDAS or α_2_δ-2 (rescue). Axonal varicosities are outlined by a dashed line. Immunolabeling for Ca_V_2.1 and synapsin (syn) revealed that, unlike α_2_δ-2, expression α_2_δ-2-ΔMIDAS in triple ko neurons rescued presynaptic synapsin but not Ca_V_2.1 clustering. The linescan patterns recorded along the indicated line support these observations. Quantification of the relative synaptic area covered by the respective immunofluorescence of presynaptic Ca_V_2.1 (**b**) and synapsin (**c**). (ANOVA with Tukey post-hoc test, ***p<0.001; Ca_V_2.1: F_(2, 88)_=27.4, p<0.001; synapsin: F_(2, 88)_=38.9, p<0.001, horizontal lines represent means, error bars SEM). Scale bar, 1 µm.

## DISCUSSION

Brain neurons simultaneously and abundantly express three different α_2_δ subunit isoforms (Cole et al., 2005; Geisler et al., 2019; Schlick et al., 2010). A fact, which, until today, has complicated studying their potentially redundant roles. By establishing a cellular α_2_δ subunit triple loss-of-function model, we here identified a critical and highly redundant role of presynaptic α_2_δ subunits in regulating glutamatergic synapse formation and differentiation, as evidenced by a series of observations: First, excitatory synapses from triple knockout cultures show a severe failure in activity dependent FM-dye uptake. Second, lack of presynaptic α_2_δ subunits strongly reduces somatic calcium currents, presynaptic clustering of the endogenous P/Q-type (Ca_V_2.1) and N-type (Ca_V_2.2) calcium channels, and the size of the active zone. Third, the failure in presynaptic differentiation is accompanied by reduced clustering of postsynaptic AMPA receptors and thinning of the postsynaptic density. Fourth, the severe synaptic phenotype can be rescued by the sole expression of α_2_δ-1, α_2_δ-2, or α_2_δ-3. Fifth, mutating the MIDAS site of α_2_δ-2 dissociates calcium channel trafficking from presynaptic synapsin clustering strongly supporting channel independent presynaptic roles of α_2_δ subunits.

### Presynaptic α_2_δ isoforms redundantly regulate synaptic differentiation of glutamatergic synapses

An increasing number of studies over the recent years have implicated calcium channel α_2_δ subunits in synaptic functions (reviewed in (Dolphin, 2018; Geisler et al., 2015). However, the severity of the phenotype of specific α_2_δ loss-of-function models strongly correlated with the expression level of the particular isoform in the affected cells or tissues: knockdown of α_2_δ-1 affected synapse formation in retinal ganglion cells (Eroglu et al., 2009), lack of α_2_δ-2 causes pre- and postsynaptic defects in hair cells of the inner ear (Fell et al., 2016), knockout of α_2_δ-3 alters presynaptic morphology of auditory nerves (Pirone et al., 2014) and in invertebrates loss-of-function of the homologous subunit resulted in abnormal presynaptic development in motoneurons (Caylor et al., 2013; Kurshan et al., 2009), and finally, the predominant expression of α_2_δ-4 in the retina (Knoflach et al., 2013) is mirrored by retinal defects and consequences on the organization of rod and cone photoreceptor synapses (Kerov et al., 2018; Wang et al., 2017; Wycisk et al., 2006). Contrary to these specialized cell types and tissues, the mammalian brain expresses all four known α_2_δ isoforms (Cole et al., 2005; van Loo et al., 2019), whereby the isoforms α_2_δ-1, −2, and −3 are strongly and most ubiquitously expressed (Geisler et al., 2019; Schlick et al., 2010). While the increasing severity of the phenotypes between α_2_δ subunit single and double knockout mice already suggested a functional redundancy, this was ultimately revealed in the cellular triple loss-of-function model established for the present study. This functional redundancy is a feature that is shared with the ubiquitous trans-synaptic adhesion proteins neuroligin and neurexin (Missler et al., 2003; Varoqueaux et al., 2006). In contrast to knockout animal models, in which the detailed cellular phenotypes may be masked by potential compensatory effects, for example by isoform redundancy or developmental adaptations, the present cellular triple knockout/knockdown model for the first time allowed analyzing the consequences of a complete lack of α_2_δ subunits in neurons from the central nervous system. Thus, our study proves that presynaptic expression of α_2_δ subunits is critical for the proper development and differentiation of excitatory glutamatergic synapses, while GABAergic synapses could still form in the absence of α_2_δ subunits. This synapse-specificity is particularly interesting as α_2_δ subunits are also critical regulators of inhibitory synapse connectivity. For example, we have recently identified that a single splice variant of the presynaptic α_2_δ-2 isoform trans-synaptically regulates postsynaptic GABA-receptor abundance and synaptic wiring (Geisler et al., 2019). Together this suggests that synapse formation and trans-synaptic signaling are two independent functions of α_2_δ subunits. The exclusive dependence of glutamatergic synaptogenesis on presynaptically expressed α_2_δ subunits is supported by the recent finding that the anti-epileptic and anti-allodynic drug gabapentin prevents synaptogenesis between sensory and spinal cord neurons by acting on presynaptic α_2_δ-1 subunits (Yu et al., 2018).

### α_2_δ subunits are critical regulators of synapse formation

In general synaptic cell adhesion molecules are thought to mediate the initial contact formation between axons and dendrites (Bury and Sabo, 2016; Garner et al., 2006). The vesicle associated protein synapsin is an early marker for presynaptic vesicle recruitment (Lee et al., 2010), yet its accumulation fails in α_2_δ triple-knockout neurons. Nevertheless, the presence of synapse-like axonal varicosities reveals an intact axo-dendritic contact formation. Together this suggests that α_2_δ subunits and therefore probably VGCC complexes take a leading role in synaptogenesis: without α_2_δ subunits excitatory synapses fail to differentiate and mutation of the MIDAS motif prevents presynaptic calcium channel trafficking but not synapsin accumulation. Previous models suggested that VGCC complexes are secondarily recruited to the release sites via their manifold interactions with presynaptic proteins (Bury and Sabo, 2016; Zamponi, 2003). Our findings also support the hypothesis that extracellular α_2_δ subunits organize the alignment of the presynaptic active zone with the postsynaptic density. Indeed, the published extracellular structure of α_2_δ-1 of the skeletal muscle Ca_V_1.1 complex (Wu et al., 2016) proposes the protrusion of α_2_δ subunits far into the synaptic cleft. Thus α_2_δ subunits may couple calcium channels with postsynaptic receptors thereby aligning the presynaptic active zone with the postsynaptic density. This hypothesis is supported by the observation that in the auditory hair cell synapse postsynaptic AMPA receptor clusters are dispersed in α_2_δ-2 knockout mice (Fell et al., 2016) and that presynaptic α_2_δ-2 regulates postsynaptic GABA-receptor abundance in GABAergic synapses (Geisler et al., 2019). Extracellular binding of α_2_δ subunits to the α_1_ subunit has been shown to be critical for efficiently coupling VGCCs to exocytosis (Hoppa et al., 2012). However, whether α_2_δ subunits interact directly or indirectly with postsynaptic receptors or trans-synaptic linkers, has yet to be elucidated. Evidently, synaptic cell adhesion molecules could provide potential candidates for such interactions. In this context it is noteworthy that α-neurexins, although being not critical for synapse formation, link presynaptic calcium channels to neurotransmitter release via extracellular domains (Missler et al., 2003; Zhang et al., 2005) and regulate presynaptic Ca_V_2.1 channels via α_2_δ subunits (Brockhaus et al., 2018). Finally, the identification of α_2_δ subunits as the first proteins that are absolutely critical for glutamatergic synapse formation paves the way for identifying up- and downstream interaction partners.

### Proposed synaptic roles for α_2_δ subunits and future implications

Taken together our study suggests an involvement of presynaptic α_2_δ subunits in several steps during synaptogenesis and synapse differentiation (Fig. 8). First, α_2_δ subunits mediate presynaptic calcium channel trafficking (Fig. 8, point 1), a role which was to be expected and which was previously demonstrated (Bauer et al., 2009). Second, α_2_δ subunits are involved in presynaptic differentiation (Fig. 8, point 2). This becomes evident by the strong effect of triple-knockout on the accumulation of synaptic vesicle-associated proteins such as synapsin and the vesicular glutamate transporter. Although it is feasible that the manifold interaction sites within the intracellular loops of calcium channel α_1_ subunits link the channel complex to synaptic vesicles (Zamponi, 2003), the partial rescue observed with the MIDAS mutation rather favors a role of α_2_δ subunits independent of the channel complex, as previously suggested (Kurshan et al., 2009). An in-depth analyses of these particular questions in our present study was impeded by the low availability of triple-knockout cultures and the necessity for pre-embedding immunolabeling with the silver-amplified gold approach to visualize the sparsely distributed, featureless boutons in electron microscopy. Thus, elucidating the precise underlying molecular mechanisms requires the future development of novel experimental tools. Third and as discussed above, α_2_δ subunits regulate postsynaptic receptor clustering and differentiation of the postsynaptic density and thus either directly or indirectly act as trans-synaptic organizers (Fig. 8, point 3a and b). As suggested by previous studies (Brockhaus et al., 2018; Geisler et al., 2019; Missler et al., 2003; Zhang et al., 2005), certain functions of α_2_δ may be modulated by their interaction with classical synaptic cell adhesion molecules.

**Fig. 8.**
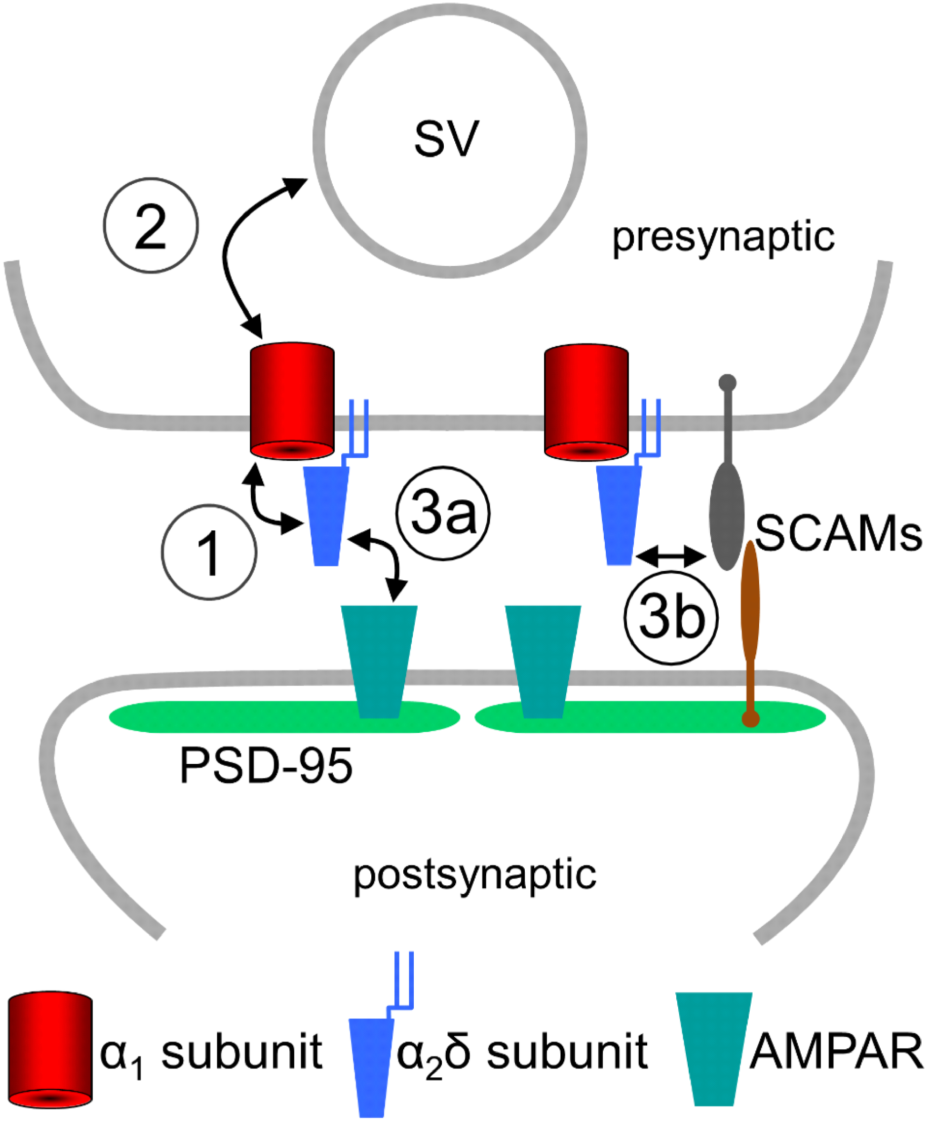
Model summarizing the putative roles of presynaptic α2δ subunits in glutamatergic synapse formation and differentiation. Our findings identified α_2_δ subunits as key organizers of glutamatergic synapses and propose their involvement in at least three critical steps during synapse maturation. By interacting with the α_1_ subunit they mediate the incorporation of VGCCs into the presynaptic active zone (1). α_2_δ subunits are involved in presynaptic differentiation and may, directly and/or indirectly via the entire VGCC complex, mediate the accumulation of synaptic vesicles (SV) to the synaptic terminal (2). Lastly, α_2_δ subunits align the presynaptic active zone with the postsynaptic membrane and postsynaptic AMPA receptors. This may be mediated by a direct interaction with AMPA receptors (3a) or by interacting with classical synaptic cell adhesion molecules (SCAMs, 3b).

Taken together, our experiments identified a critical role of presynaptic α_2_δ subunits in glutamatergic synapse differentiation. This affects our current view on excitatory synapse formation and implicates α_2_δ subunits and therefore presynaptic calcium channel complexes as potential nucleation points for the organization of synapses.

## ACKNOWLEDGEMENTS

We thank Arnold Schwartz for providing α_2_δ-1 knockout mice, Ariane Benedetti and Sabine Baumgartner for technical support, Daniel Gütl from the Electron Microscopy Facility at ISTA for help with experiments, Hermann Dietrich, Anja Beierfuß and her team for animal care, Jutta Engel and Jörg Striessnig for critical discussions, and Bruno Benedetti and Bernhard Flucher for critical discussions and reading the manuscript. This study was supported by the Austrian Science Fund (FWF) grants P24079, F44060, F44150, DOC30-B30 (G.J.O) and T855 (M.C.), and from European Research Council (ERC) grant AdG 694539 (R.S.). This work is part of the PhD theses of C.L.S. and S.G.

## AUTHOR CONTRIBUTIONS

C.L.S. designed, performed and analyzed experiments and wrote the manuscript, B.S. and B.N. participated in the experimental work and data analysis, R.I.S., S.G., W.A.K., and R.S. designed, performed and analyzed experiments, M.C. planned and performed molecular biological experiments, G.J.O. conceived the study, supervised the project, designed and analyzed experiments and wrote the manuscript. All authors discussed, modified and approved the final manuscript.

## COMPETING FINANCTIAL INTERESTS

The authors declare that they have no competing interests.

## SUPPLEMENTARY INFORMATION

**Suppl. Fig. 1.**
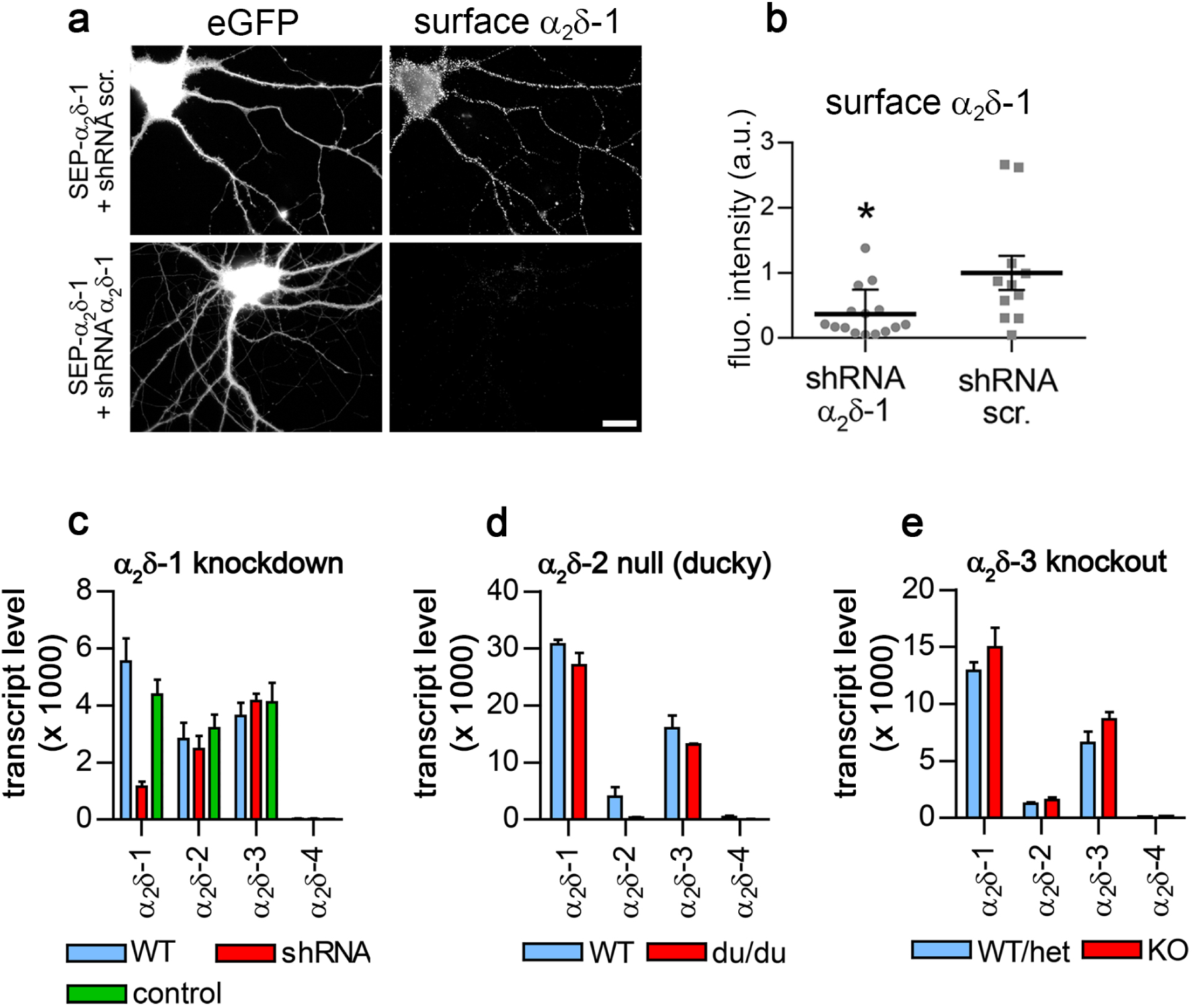
shRNA knockdown of α_2_δ-1 protein and mRNA in cultured hippocampal neurons. (**a**) Live cell surface staining of neurons expressing α_2_δ-1 with an extracellular super-ecliptic pHluorin tag (SEP-α_2_δ-1) reveals robust knockdown of protein expression by α_2_δ-1 specific shRNA (lower panel) when compared to scrambled control shRNA. Scale bar, 20 μm. (**b**) shRNA decreased surface expression of SEP-α_2_δ-1 to 37±10% (mean±SD) of control neurons transfected with scrambled shRNA [t_(12)_=2.3, p=0.044]. (**c-e**) qPCR expression profiles of the four α_2_δ isoforms in three different α_2_δ deficient model systems. (**c**) Lentiviral transfection of cultured hippocampal neurons (DIV 24; ∼90% transfection efficiency) with α_2_δ-1 specific shRNA (shRNA; red bars) significantly reduced α_2_δ-1 transcript levels to 25±3% (p=0.015) of neurons transfected with scrambled control shRNA (green bars) and to 21±6% (p=0.005) of untransfected neurons (WT; blue bars; ANOVA with Holm-Sidak posthoc test). Loss of either α_2_δ-2 in ducky mice (**d**, α_2_δ-2 null) or α_2_δ-3 (**e**, α_2_δ-3 knockout mice) did not induce compensational changes in the expression levels of the other isoforms. Mutated α_2_δ-2 mRNA in ducky mice is unstable and thus strongly reduced, whereas α_2_δ-3 mRNA with the Lac-Z insert is stably expressed. [n-numbers: α_2_δ-1, 3 culture preparations; ducky, 2 hippocampus preparations from 8 week old mice; α_2_δ-3 knockout, 3 hippocampus preparations from 8 week old mice]

**Suppl. Fig. 2.**
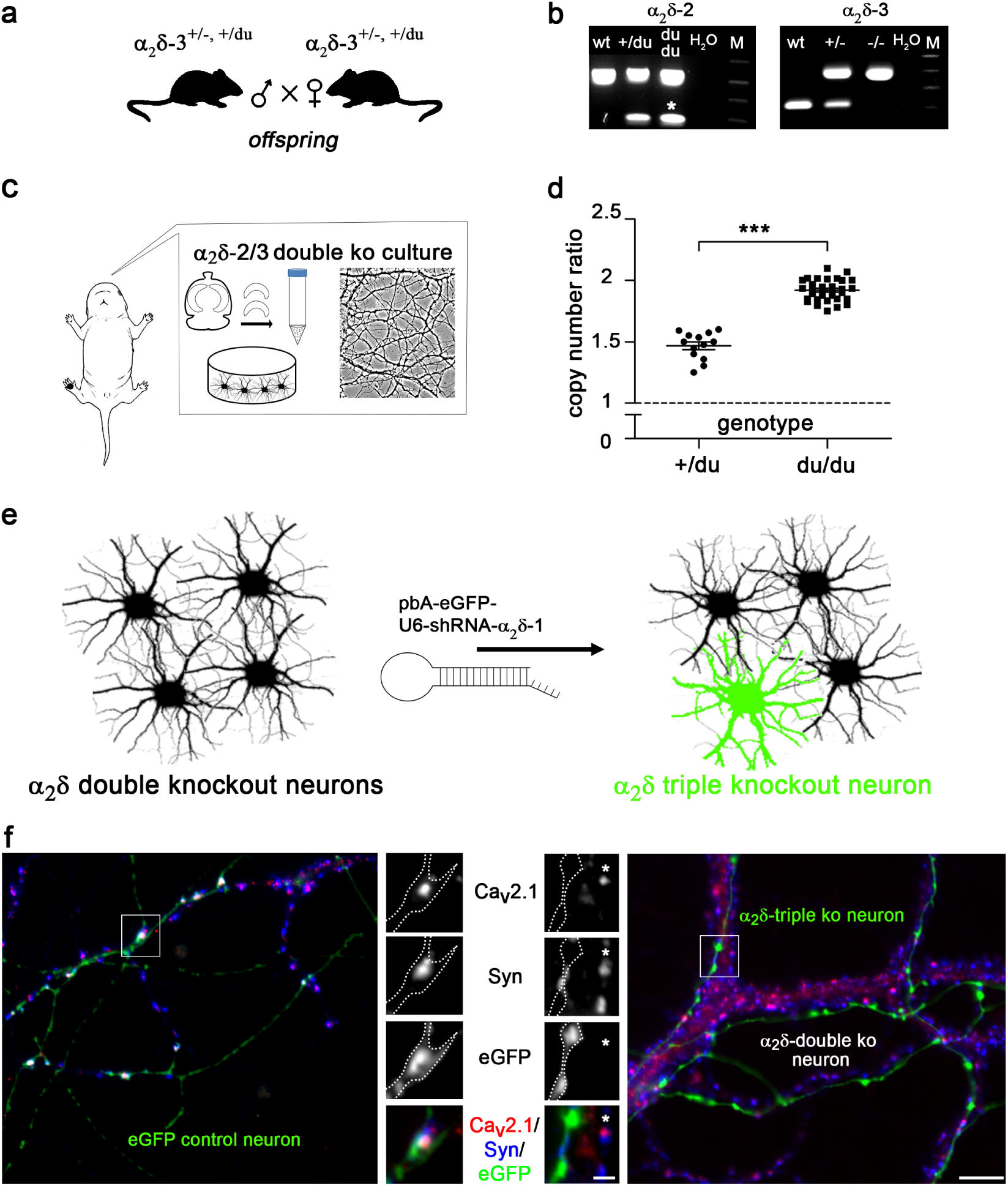
Establishing a cellular α2δ subunit triple knockout/knockdown model. (**a**) Crossbreeding double heterozygote α_2_δ-2^+/du^ / α_2_δ-3^+/-^mice yielded α_2_δ-2/-3 double knockout pups with a probability of 6.25 %. (**b**) On P0 neonatal pups were individually marked by paw-tattooing. The α_2_δ-2 wildtype allele was identified by a 541 bp band (wt). The presence of the du allele in heterozygote (+/du) and homozygote ducky (du/du) mice was indicated by an additional band at ∼280 bp (see Materials and Methods for details), whereby homozygosity (du/du) was suggested by an increased intensity of the ∼280 bp band (asterisk). Genotyping for α_2_δ-3 revealed bands at 183 bp for wt, 331 and 183 bp for heterozygote, and 331 bp for homozygote knockout mice. (**c**) Putative double knockout pups (α_2_δ-3^-/-,^ α_2_δ-2^du/du^) were selected for hippocampal culture preparation in parallel with control littermates. (**d**) Due to the large genomic rearrangement in ducky mice, the ducky mutation required a final confirmation employing a copy number counting qPCR approach. The graph indicates experimentally determined copy number ratios from 32 (wt and du/du) and 13 (+/du) mice: wt (α_2_δ-2^+/+^), 2 copies normalized to a ratio of 1 (dashed line); heterozygote (α_2_δ-2^+/du^), 3 copies a ratio of ∼ 1.5; homozygote (α_2_δ-2^du/du^), 4 copies a ratio of ∼2.0. (ANOVA on ranks with Dunn’s post-hoc test, ***p<0.001). (**e**) Ultimately, in confirmed α_2_δ-2/-3 double knockout cultured hippocampal neurons α_2_δ triple knockout neurons were established by transfection with α_2_δ-1 shRNA (light green). (**f**) **left**: Hippocampal neurons from control litters transfected with eGFP show axonal varicosities (eGFP), positively labeled for synapsin and the presynaptic Ca_V_2.1 calcium channel. **right:** Axons from eGFP positive triple knockout neurons displayed varicosities similar to control neurons. However, the majority of these varicosities lacked staining for synapsin and Ca_V_2.1. Importantly, axons and varicosities from triple knockout neurons can be directly compared to eGFP negative double knockout synapses (on the same postsynaptic dendrites, red/Ca_V_2.1, blue/synapsin; asterisks in the magnified selection). Scale bars, 1µm and 5µm.

**Suppl. Fig. 3.**
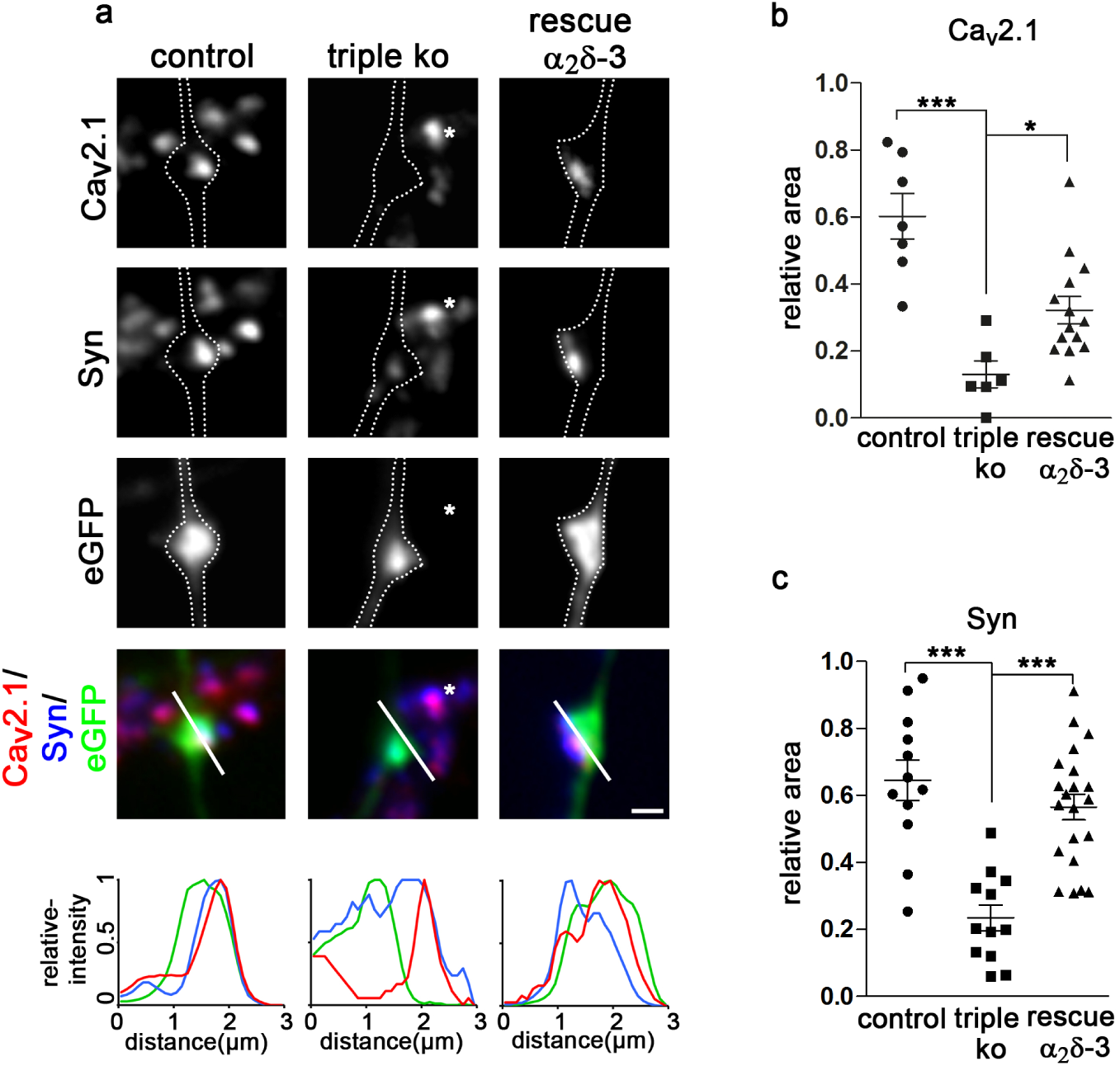
The role of α2δ subunits in synapse formation is highly redundant. (**a**) Immunofluoresence analysis of axonal varicosities from wildtype neurons (control, neurons transfected with eGFP only), triple knockout neurons (triple ko, α_2_δ-2/-3 double knockout neurons transfected with shRNA-α_2_δ-1 plus eGFP), and triple knockout neurons expressing α_2_δ-3 (rescue, α_2_δ-2/-3 double knockout neurons transfected with shRNA-α_2_δ-1 plus eGFP and α_2_δ-3). Putative presynaptic boutons were identified as eGFP-filled axonal varicosities along dendrites of untransfected neurons (confer Suppl. Fig. 2) and outlined with a dashed line. Outlines indicate synaptic boutons from control, triple ko as well as α_2_δ-3 rescued neurons. The failure in synapse formation is indicated by the highly reduced area of synapsin as well as presynaptic Ca_v_2.1. Interestingly this defect could be rescued by each individual α_2_δ-isoform (shown is α_2_δ-3). The redundant function of α_2_δ subunits in synapse formation and calcium channel targeting is further supported by the fact that neighboring eGFP-negative α_2_δ-2/-3 double ko boutons still containing α_2_δ-1 formed synapses together with proper Ca_v_2.1 abundance (asterisks). (**b, c**) Quantification of the fluorescence intensities of presynaptic Ca_V_2.1 and synapsin clustering in control, triple knockout, and α_2_δ-3-expressing (rescued) triple knockout neurons (ANOVA with Holm-Sidak post-hoc test, *p<0.05, ***p<0.001; Ca_V_2.1: 6-14 cells from 2-3 culture preparations; Synapsin: 12-17, 4). Error bars indicate SEM. Scale bar, 1µm.

**Supplementary Table 1.** Effect of α2δ knockout on properties of total endogenous calcium channels.

| | control | | | triple ko | | | rescue $\alpha_2\delta$-2 | | |
| --- | --- | --- | --- | --- | --- | --- | --- | --- | --- |
| | Mean | ±SEM | n | Mean | ±SEM | n | Mean | ±SEM | n |
| CD (pA/pF) | -75.3 | 5.8 | 14 | -32 | 3.6 | 10 | -71.3 | 6.6 | 8 |
| $V_{50\text{act}}$(mV) | -17.1 | 2.2 | 14 | -16.9 | 2.2 | 10 | -14.1 | 1.9 | 8 |
| $V_{\text{rev}}$ (mV) | 48.4 | 0.6 | 14 | 34.6 | 2.4 | 10 | 46.4 | 1.7 | 8 |

| | control | | | $\alpha_2\delta$-3 ko | | | $\alpha_2\delta$-2/3 ko | | |
| --- | --- | --- | --- | --- | --- | --- | --- | --- | --- |
| | Mean | ±SEM | n | Mean | ±SEM | n | Mean | ±SEM | n |
| CD (pA/pF) | -75.3 | 5.8 | 14 | -66.3 | 4.2 | 18 | -51.6 | 3.5 | 26 |
| $V_{50\text{act}}$(mV) | -17.1 | 2.2 | 14 | -14.6 | 1.8 | 18 | -18 | 1.8 | 26 |
| $V_{\text{rev}}$ (mV) | 48.4 | 0.6 | 14 | 46.7 | 1.0 | 18 | 44.7 | 1.4 | 26 |

CD, current density; $V_{50\text{act}}$ , half-maximal voltage of activation; $V_{\text{rev}}$ , reversal potential.

